# Design, Synthesis and Evaluation of Inhibitors of the SARS-CoV2 nsp3 Macrodomain

**DOI:** 10.1101/2022.02.27.482176

**Authors:** Lavinia M. Sherrill, Elva E. Joya, AnnMarie Walker, Anuradha Roy, Yousef M. Alhammad, Moriama Atobatele, Sarah Wazir, George Abbas, Patrick Keane, Junlin Zhuo, Anthony K.L. Leung, David K. Johnson, Lari Lehtiö, Anthony R. Fehr, Dana Ferraris

## Abstract

A series of amino acid based *7H*-pyrrolo[2,3-*d*]pyrimidines were designed and synthesized to discern the structure activity relationships against the SARS-CoV-2 nsp3 macrodomain (Mac1), an ADP-ribosylhydrolase that is critical for coronavirus replication and pathogenesis. Structure activity studies identified compound **15c** as a low-micromolar inhibitor of Mac1 in two ADP-ribose binding assays. This compound also demonstrated inhibition in an enzymatic assay of Mac1 and displayed a thermal shift comparable to ADPr in the melting temperature of Mac1 supporting binding to the target protein. A structural model reproducibly predicted a binding mode where the pyrrolo pyrimidine forms a hydrogen bonding network with Asp^22^ and the amide backbone NH of Ile^23^ in the adenosine binding pocket and the carboxylate forms hydrogen bonds to the amide backbone of Phe^157^ and Asp^156^, part of the oxyanion subsite of Mac1. Compound **15c** also demonstrated notable selectivity for coronavirus macrodomains when tested against a panel of ADP-ribose binding proteins. Together, this study identified several low MW, low μM Mac1 inhibitors to use as small molecule chemical probes for this potential anti-viral target and offers starting points for further optimization.

**Graphical Abstract:** 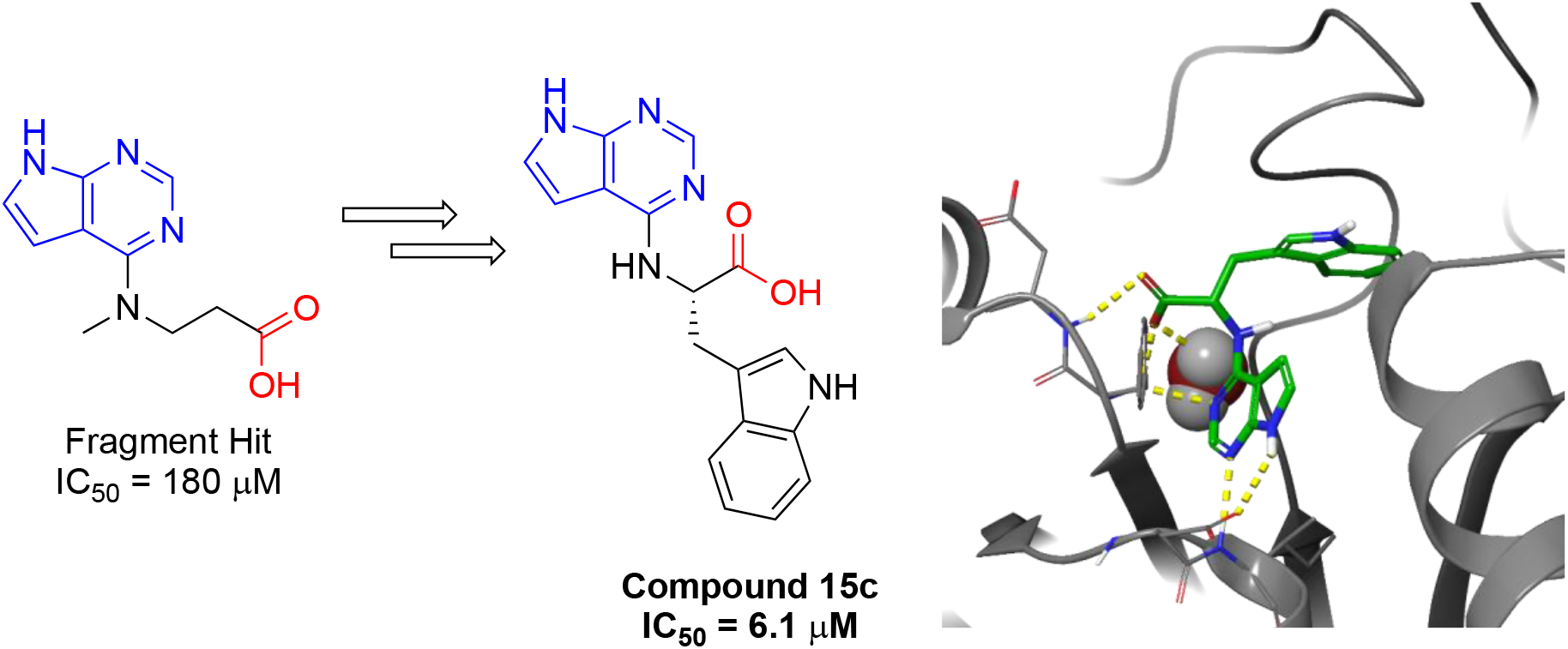

## Introduction

The current outbreak of SARS-CoV-2 and other recent outbreaks of highly pathogenic human coronaviruses (CoVs) has exposed the challenges posed by CoVs and the lack of antivirals available to target CoVs and other viruses of pandemic potential. The development of novel antivirals targeting conserved CoV proteins that could counter emerging CoVs is greatly needed, and many different approaches are currently being pursued [1]. One potential novel CoV drug target is the highly-conserved macrodomain (Mac1), which is a domain within the large non-structural protein 3 (nsp3) protein. Mac1 is present in all CoVs, and contains a conserved three-layered α/β/α fold, typical of all macrodomains. All CoV Mac1 proteins tested have ADP-ribose binding (reader) and ADP-ribosylhydrolase (ARH) (eraser) activity and are members of the larger MacroD-type macrodomain family, which includes human macrodomains Mdo1 and Mdo2 [2]. Importantly, several studies have demonstrated that Mac1 is essential for CoV-mediated pathogenesis in multiple animal models of infection, indicating that Mac1 may be a suitable drug target [3].

ADP-ribosylation is a dynamic post-translational modification catalyzed by ADP-ribosyltransferases (ARTs, a.k.a. Sirtuins, ARTCs and PARPs) where ADP-ribose is transferred from NAD^+^ onto target proteins or to DNA/RNA [4]. ADP-ribose can be transferred as a mono-ADP-ribose (MAR), or consecutively attached units of MAR covalently attached through glycosidic bonds to preceding ADP-ribose units to form a branched or linear poly-ADP-ribose (PAR) chain. ADP-ribose can then be removed from proteins through several different eraser enzymes, such as PAR glycohydrolase (PARG), ADP-Ribosylhydrolases (ARHs), and macrodomains. Although these proteins catalyze the same reaction, the families are structurally distinct [2]. ADP-ribosylation is known to play important biological roles such as the DNA damage response, cellular signaling, cell cycle control, ER stress, immune regulation, and many others. Furthermore, both mono- and poly-ARTs can inhibit virus replication, implicating ADP-ribosylation in the host innate immune response to infection [5]. Furthermore, in addition to CoVs, several families of viruses encode for macrodomains including the *Coronaviridae, Togaviridae, Matonaviridae, Hepeviridae*, and the *Iridoviridae*, indicating a significant evolutionary advantage for some these viruses to directly counter ADP-ribosylation.

Multiple studies have reported that Mac1 is critical for CoV replication and pathogenesis. This has largely been accomplished using reverse genetics to mutate a highly conserved asparagine to alanine (N41A-SARS-CoV). This mutation nearly eliminates the enzymatic activity of a recombinant SARS-CoV macrodomain protein *in vitro* [6]. This mutation had only a limited impact on CoV replication in transformed cells, but in animal models these mutant viruses proved highly attenuated, with low viral loads, increased IFN production, and the inability to cause significant disease [6–10]. Recombinant murine hepatitis virus strain JHM (MHV-JHM) with this same mutation (N1347A) replicated poorly in primary macrophages, but importantly this defect could be partially rescued by the PARP inhibitors or siRNA knockdown of PARP12 or PARP14 [11]. These data indicate that Mac1 likely counters PARP-mediated anti-viral ADP-ribosylation. Further genetic analysis found that mutations in MHV Mac1 predicted to alter ADP-ribose binding resulted in severe replication defects in cell culture (D1329A) or could not be recovered (G1439V, D1329A/N1347A), indicating that for some CoVs, Mac1 may be essential for replication [12]. Looking beyond CoVs, mutations in Chikungunya virus, Sindbis virus, and Hepatitis E virus macrodomains also have severe phenotypic effects on virus replication and pathogenesis [13–17]. As viral macrodomains are clearly important virulence factors, they are potential targets for anti-viral therapeutics [3].

Since the emergence of SARS-CoV-2, there has been an increased effort to identify compounds that bind to or inhibit the activity of the SARS-CoV-2 Mac1 protein. Most of these studies only involved molecular modeling of potential inhibitors, though a few have tested compound activity in biochemical assays [18–22]. Virdi et al. used a differential scanning fluorescence (DSF) assay to screen ~2500 compounds, and while they identified several compounds that altered the melting temperature of the protein, it’s yet unclear if any of these compounds can inhibit Mac1 activity [21]. Russo et al. developed an immunofluorescence assay to measure Mac1 activity in cells. They demonstrated that poly(I:C) or IFNγ induced global MARylation in A549 cells that could be removed by expression of Mac1, but not the N1040A mutant protein that has minimal enzymatic activity [19]. They utilized this assay in a small inhibitor screen but did not identify any significant hits. Despite this, the assay could be a highly useful tool for measuring compound activity in cells. Dasovich et al. developed a novel ADP-ribosylhydrolase assay that utilizes the ability of NudF to specifically cleave AMP from ADP-ribose released from ADP-ribosylated proteins in the presence of macrodomain, but not from untreated ADP-ribosyalted protein [18]. The generated AMP can then be measured in high-throughput fashion. The authors screened over 3000 compounds using this assay and identified 2 compounds that inhibited mono-ARH activity. One, Dihydralazine, inhibited both Mdo2 and the SARS-CoV Mac1 domain with high IC_50_ values (~0.5 mM), while the other compound, Dasatinib, specifically inhibited CoV Mac1 mono-ARH activity with an IC_50_ of ~50 μM. While Dasatinib is toxic to cells at higher concentrations, it may be a suitable scaffold for further inhibitor development. A FRET assay used in the present publication was tested by screening of approved drugs and suramin was found to be an inhibitor of Mac1 with an IC_50_ of ~8.7 μM, and it also showed concentration dependent stabilization of the protein in DSF assay. Suramin has however multiple reported activities and does not therefore represent a high quality chemical probe for Mac1 inhibition [22].

Recently, a screen was conducted in a multi-national collaboration to identify several small molecule fragments as weak inhibitors of the SARS-CoV2 nsp3 macrodomain [20]. One such inhibitor, a fragment with a pyrrolo pyrimidine core (blue heterocycle, Figure 1), compound **1**, is shown in Figure 1 below. This compound demonstrated weak potency against the macrodomain (IC_50_ = 180 μM). In addition, the crystal structure of this compound in the active site of the nsp3 macrodomain demonstrated a hydrogen bonding network between the pyrrolo pyrimidine and the side chain of Asp^22^ and the amide backbone NH of Ile^23^ in the adenosine binding site of the nsp3 macrodomain (Figure 1) [20]. In addition, one crystal structure showed the side chain carboxylate of **1** (red, Figure 1) forming hydrogen bonds with the backbone amides of Phe^157^ and Asp^156^, part of the oxyanion subsite. This pyrrolo pyrimidine core was also part of another fragment, compound **2**, (IC_50_ = 400 μM), with the same binding mode. Taken together, these data provide a good starting point for optimization for several reasons: 1) the molecular weights of these two fragment hits are <300, allowing for fragment growth strategies that will leave the final inhibitors within drug like parameters; 2) the pyrrolo pyrimidine core was the most common core found among the hits and it has a conserved/predictable binding mode; 3) proximity to oxyanion subsite and phosphate subsites to further block ADPr binding/recognition and improve potency; 4) derivatives with this core can be synthesized readily from commercially available starting materials.

**Figure 1.**
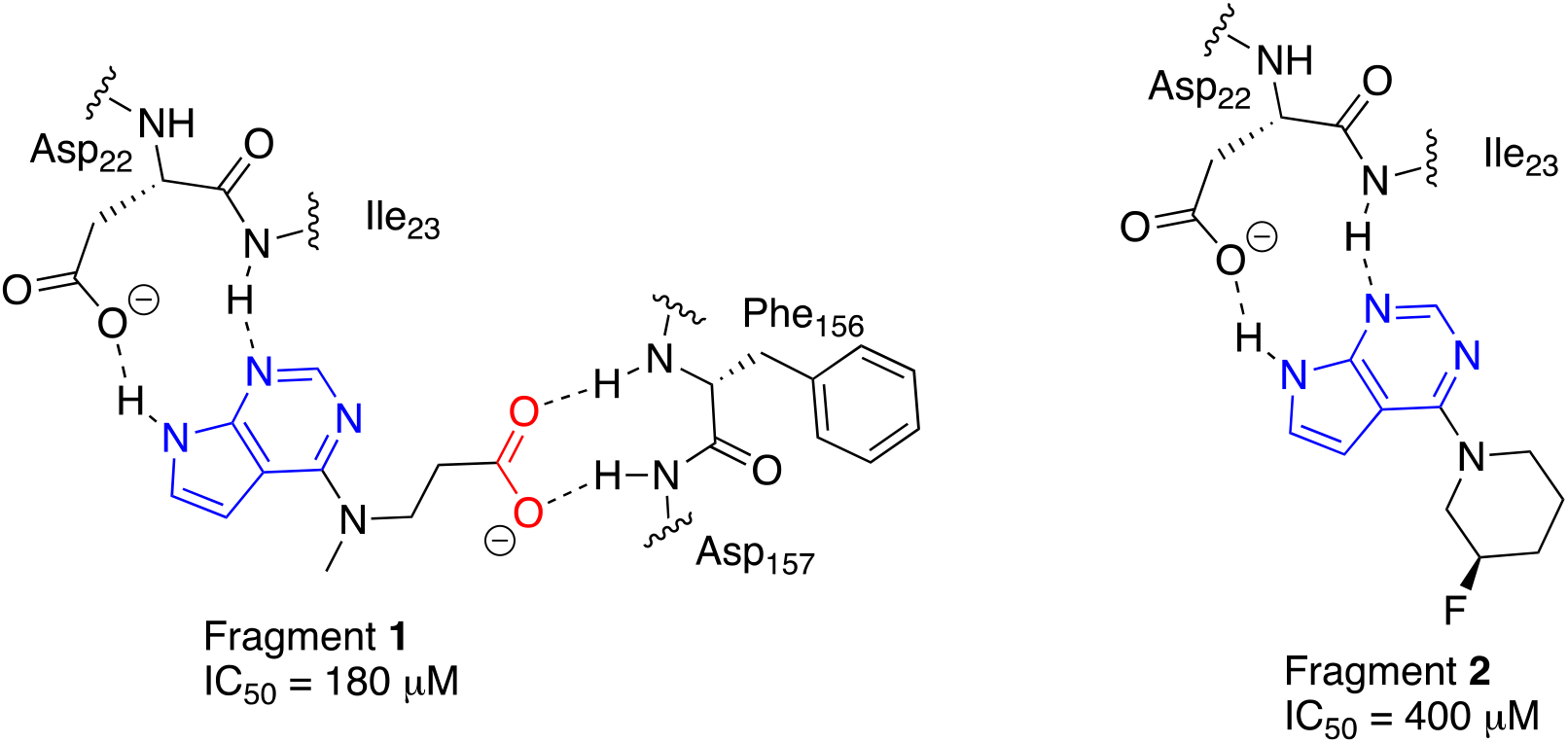
Binding mode for indolo pyrimidine fragment hits generated from co-crystal structures of 1 (PDB id. 5RSG) and 2 (PDB id. 5RSE) with Mac1.

This manuscript outlines the design, synthesis and *in vitro* evaluation of two subseries of pyrrolo pyrimidines. The subseries derived from secondary amino acids identified two compounds **6g** and **6h** that improved the potency almost 10-fold over the initial fragment. The subseries derived from primary amino acids afforded compound **15c**, a low micromolar derivative (IC_50_ = 6.1 μM) capable of causing a 4.3 °C thermal shift in the DSF assay, comparable to that of ADPr, consistent with direct binding to Mac1. Compound **15c** also demonstrated the ability to inhibit enzymatic activity of Mac1. Molecular modelling allowed us to rationalize the potency observed for the derivatives as well as a rationale for the improvement in potency from compound **15c** through fragment growth. Furthermore, compound **15c** was selective for coronavirus macrodomains as indicated by profiling against a panel of ADP-ribose binding proteins. Taken together, these two pyrrolo pyrimidine series yielded compounds with improved potency capable of selectively inhibiting Mac1 ADP-ribose binding and ARH activity.

## Chemistry

Initial designs for inhibitors focused on attaching a carboxylic acid 1-3 atoms away from the pyrrolo pyrimidine core, similar to that of compound **1**. The general synthesis of secondary cyclic amino acid derivatives is outlined in Scheme 1 below. Commercially available chloride **3** was the starting material for a nucleophilic aromatic substitution reaction with secondary amino esters **4a-b** and **4f-h**. This reaction was conducted with K_2_CO_3_ in DMF or with DIEA in isopropanol to afford the desired esters **5a-b** and **5f-h** in good yield. Ester hydrolysis in either acidic or basic conditions led to the production of carboxylic acids **6a-b** and **6f-h**. Alternatively, secondary amino acids **7a-c** reacted with chloride **3** to afford the desired acids **6c-e** directly.

**Scheme 1.**
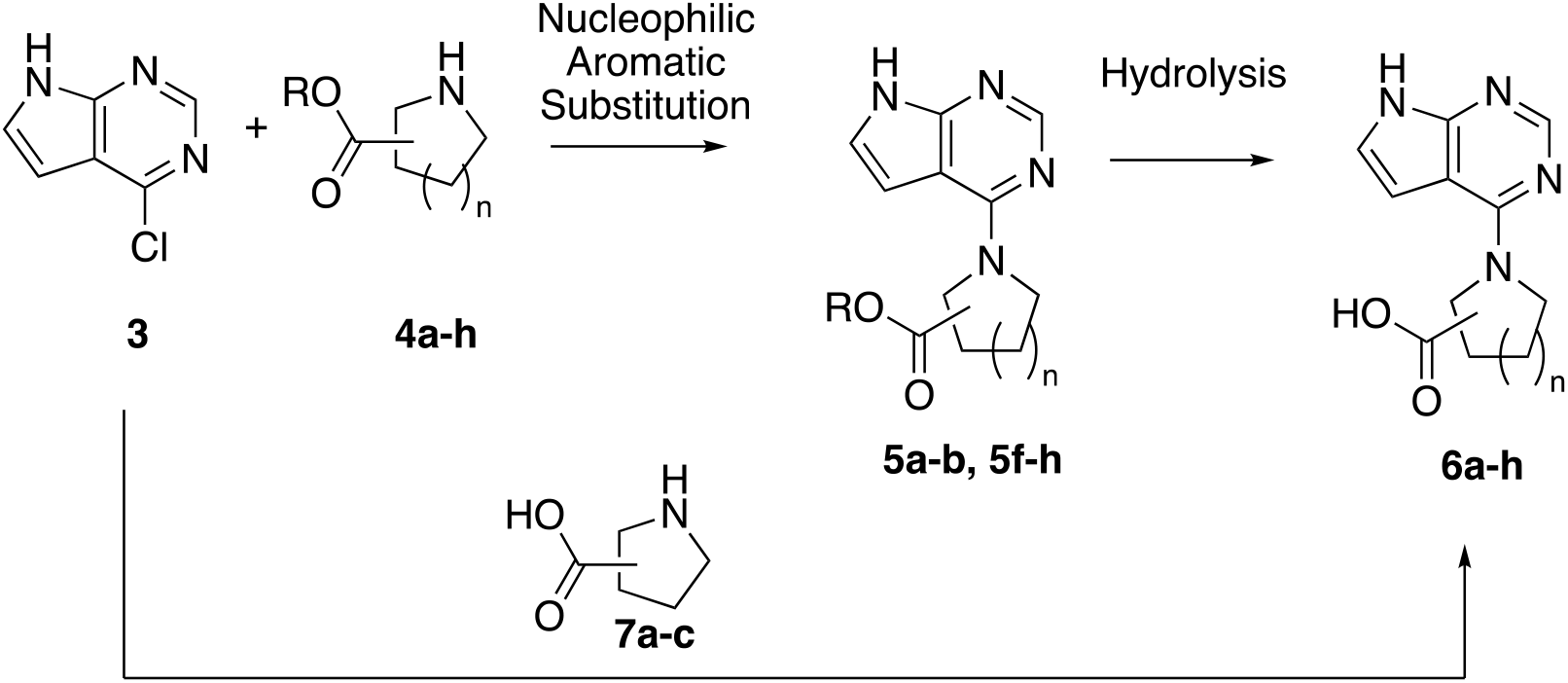
Synthesis of secondary amino acid derivatives.

The next series of compounds focused on incorporating the carboxylic acid into a piperazine ring. The synthesis of this series is shown in Scheme 2. Nucleophilic aromatic substitution of chloride **3** with (*R*)- and (*S*)-piperazine esters **8a-b** afforded **9a-b** in good yield. Deprotection of the Boc group from **9a-b** using TFA afforded esters **10a-b** as TFA salts. Basic hydrolysis of esters **9a-b** led to the free carboxylates **11a-b**. Deprotection of the boc group from carboxylates **11a-b** afforded the TFA salts of the amino acids **12a-b** in good yield.

**Scheme 2.**
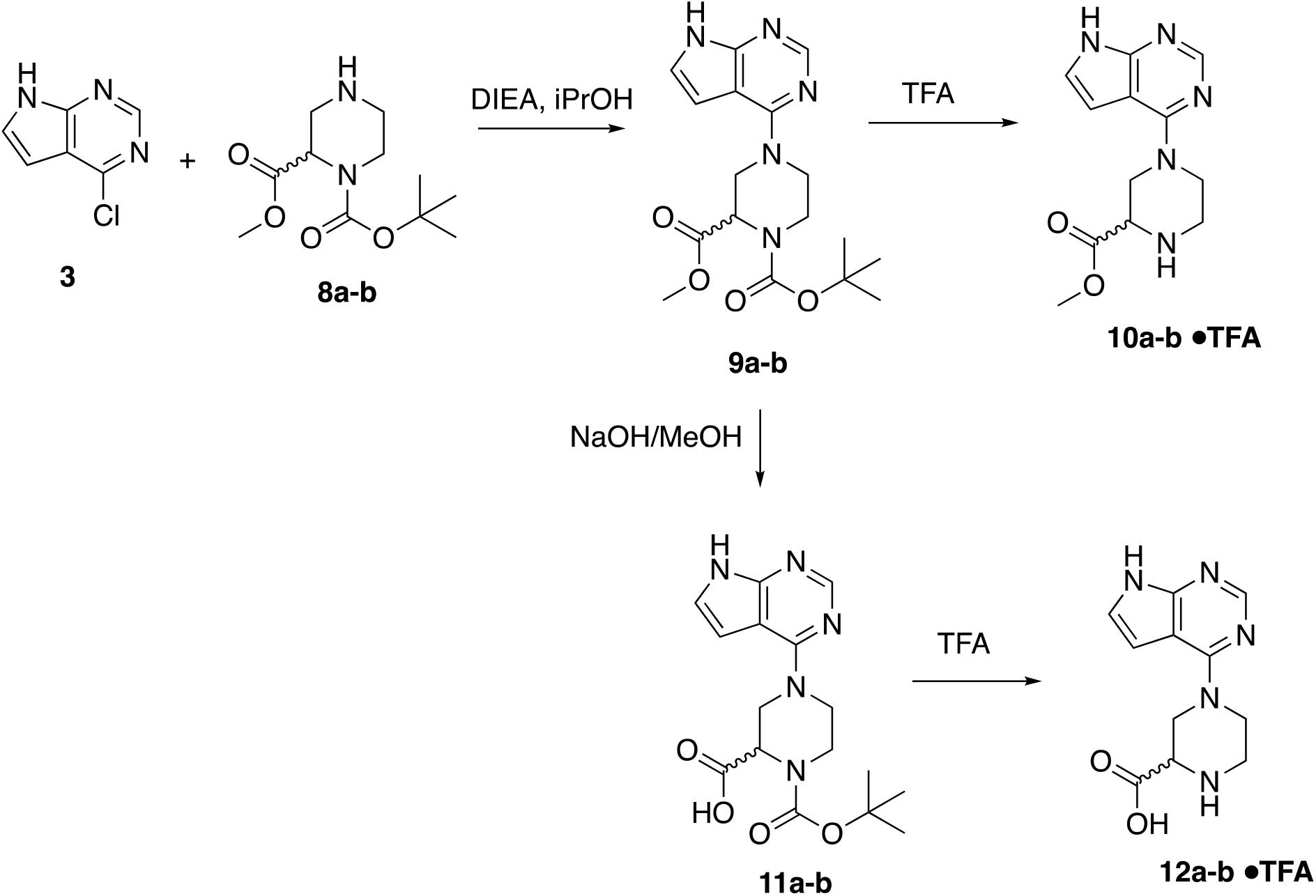
Synthesis of piperazine based pyrrolo pyrimidines.

Synthesis of the natural and non-natural primary amino acid pyrrolo pyrimidines is outlined in Scheme 3 below. The nucleophilic aromatic substitution was conducted with either the amino esters **13a-c** or in the case of glycine, with the amino acid itself to form **15d**. Hydrolysis of the esters **14a-c** afforded the desired amino acid pyrrolo pyrimidines **15a-c**.

**Scheme 3.**
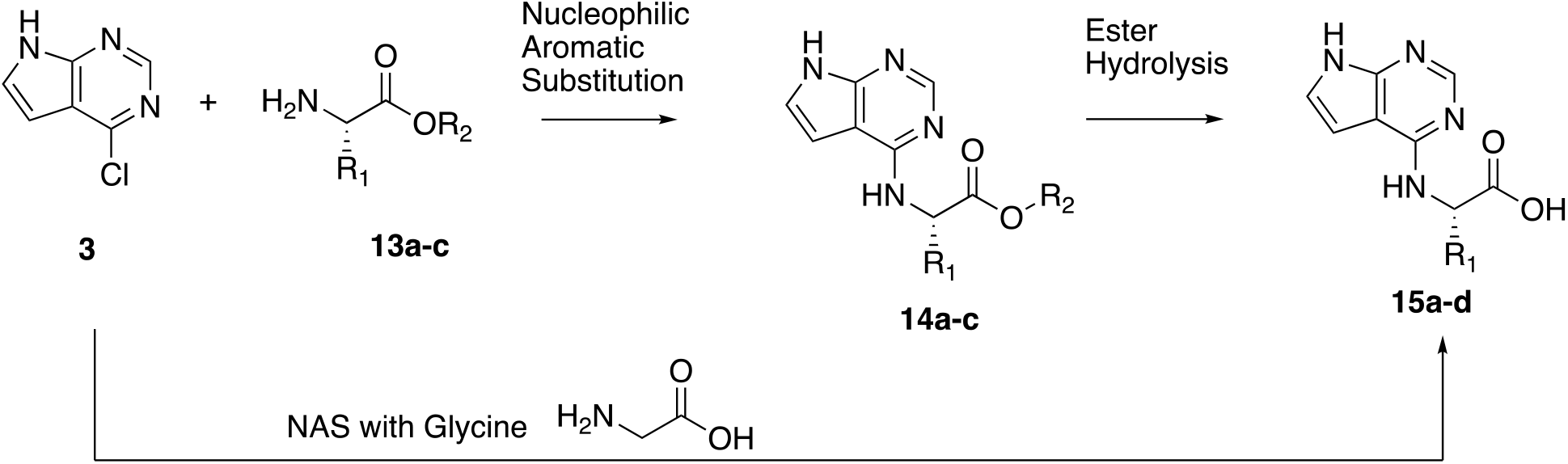
Synthesis of primary natural amino acid derivatives.

The beta-amino acid derivative was synthesized as outlined in Scheme 4 below.

**Scheme 4.**
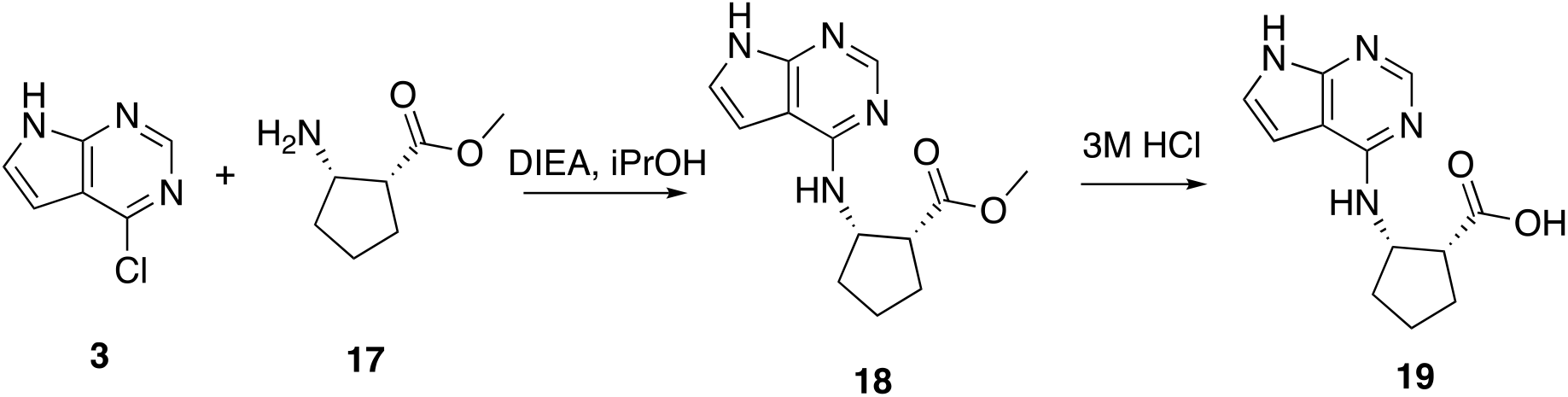
Synthesis of primary beta amino acid derivative **19**

Nucleophilic aromatic substitution of amino ester **17** with chloride **3** afforded the amino ester **18** in moderate yield. Hydrolysis to the carboxylic acid was accomplished in 3M HCl to afford the desired carboxylate **19** in 63% yield.

## Results and Discussion

Preliminary binding data of fragments **1** and **2** indicated that the pyrrolo pyrimidine core forms a tight hydrogen bond network with Asp^22^, Ile^23^ and in some cases Phe^156^ [20]. Therefore, we maintained these three features while modifying the 4-position on this heterocyclic ring. Our first round of derivatization attempted to take advantage of two more hydrogen bonds that can potentially be formed by interactions with the backbone of Phe^156^ and Asp^157^ and a carboxylic acid moiety from the inhibitor (Figure 1). We hypothesized that this type of interaction would improve the potency by taking advantage of the entropic gains of constraining the carboxylic acid within a ring approximately 1-3 atoms away from the pyrrolo pyrimidine core. Based on the binding of compound **2**, it is likely that rings of approximately 4-6 atoms will be tolerated within the adenosine binding site, and indeed this is what the data from the structure activity studies indicate (Table 1). To assess the ability for binding, we used a previously published AlphaScreen™ (amplified luminescence proximity homogeneous assay) (AS) displacement binding assay and adapted it for SARS-CoV-2 Mac1 [23]. This assay uses a mono-ADP-ribosylated and biotinylated peptide with a His6-tagged SARS-CoV-2 or Mdo2 macrodomain. Importantly, the ADP-ribose is attached via an aminoxy bond, essentially eliminating the chance that the enzymatically active macrodomains would not cleave it from the peptide. The general SAR trend that was noticed from the cyclic amino esters (**5a-b, d, f-h**) and amino acid derivatives (**6a-h**) was that the ester derivatives were all less potent than their corresponding carboxylic acid derivatives as shown by comparing **5b/6b**, **5f/6f**, **5g/6g** and **5h/6h**. Often times the carboxylic acid is several fold more potent than the ester as seen between **5g** (IC_50_ = 127μM) and **6g** (IC_50_ = 21.6μM). This trend supports our hypothesis that the carboxylic acid is important for additional binding. Another trend that was noted within this series was that carboxylic acids incorporated into 6 membered rings tended to afford compounds with greater potency. The two most potent compounds in this series, **6g** and **6h** were both derived from 6-membered piperidines and were several-fold more potent than any carboxylate from either a 4-membered azetidine (**6a-b**) or a 5-membered pyrrolidine (**6c-f**). In addition, **6g** and **6h** demonstrated selectivity for the SARS-CoV-2 macrodomain exhibiting minimal inhibition of the human macrodomain Mdo2 (>300μM).

**Table 1.**
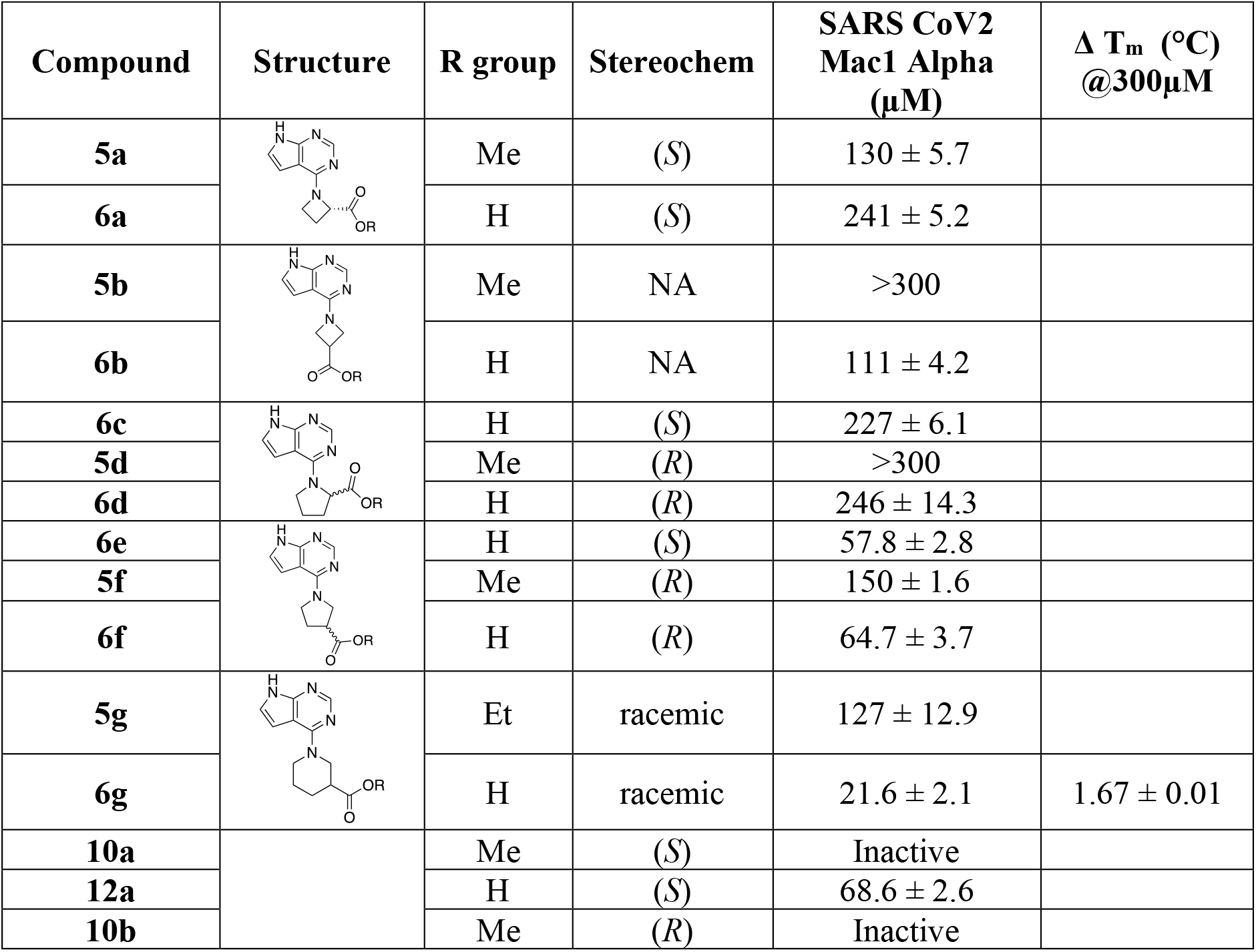
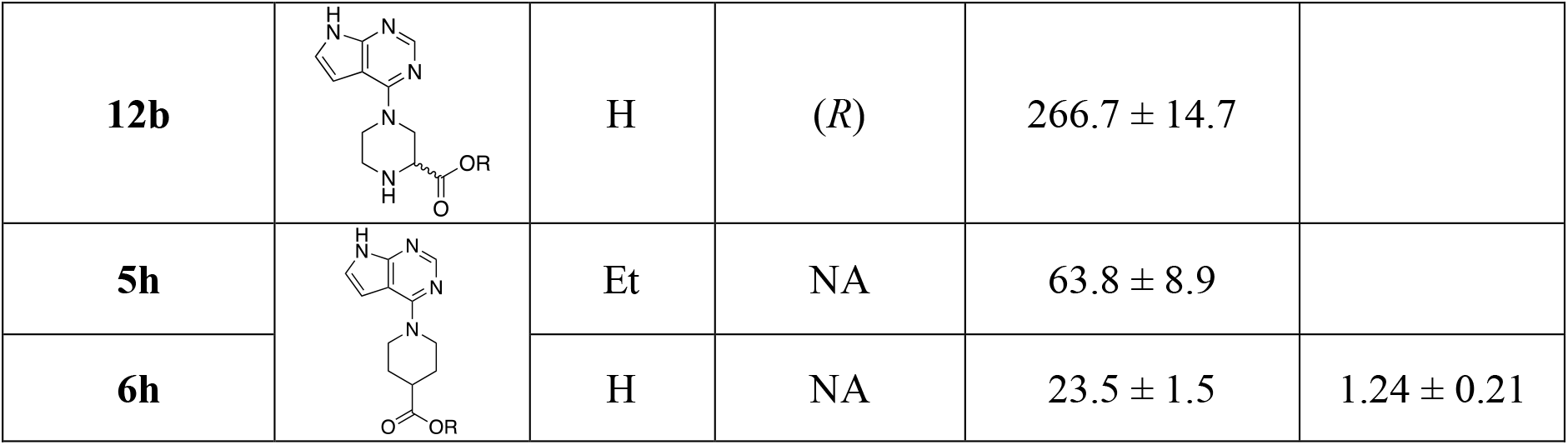
Mac1 Inhibition data for secondary amino acid derivatives. IC_50_s, changes in the melting temperature and corresponding SDs of three replicates are given.

| Compound | Structure | R group | Stereochem | SARS CoV2 Mac1 Alpha ($\mu M$) | $\Delta T_m$ ($^{\circ}C$) @300$\mu M$ |
| --- | --- | --- | --- | --- | --- |
| **5a** | | Me | (*S*) | $130 \pm 5.7$ | |
| **6a** | | H | (*S*) | $241 \pm 5.2$ | |
| **5b** | | Me | NA | $>300$ | |
| **6b** | | H | NA | $111 \pm 4.2$ | |
| **6c** | | H | (*S*) | $227 \pm 6.1$ | |
| **5d** | | Me | (*R*) | $>300$ | |
| **6d** | | H | (*R*) | $246 \pm 14.3$ | |
| **6e** | | H | (*S*) | $57.8 \pm 2.8$ | |
| **5f** | | Me | (*R*) | $150 \pm 1.6$ | |
| **6f** | | H | (*R*) | $64.7 \pm 3.7$ | |
| **5g** | | Et | racemic | $127 \pm 12.9$ | |
| **6g** | | H | racemic | $21.6 \pm 2.1$ | $1.67 \pm 0.01$ |
| **10a** |  | Me | (*S*) | Inactive |  |
| **12a** | | H | (*S*) | $68.6 \pm 2.6$ | |
| **10b** |  | Me | (*R*) | Inactive |  |
| **12b** | | H | (*R*) | $266.7 \pm 14.7$ | |
| **5h** | | Et | NA | $63.8 \pm 8.9$ | |
| **6h** | | H | NA | $23.5 \pm 1.5$ | $1.24 \pm 0.21$ |

The piperazine derivatives shown in Scheme 2 were initially synthesized to provide another potential site for fragment expansion, namely the nitrogen atom not connected to the pyrrolo pyrimidine. Unfortunately, neither enantiomer of the methyl esters (**10a-b**) nor the carboxylic acids (**12a-b**) were as potent as the piperidine derivatives **6g** and **6h**.

Molecular modeling was performed for compounds **6g** and **6h** as shown in Figure 2. The co-crystal structures with parent compound **1** and ADPr have previously been published [20, 24] and are shown (Figure 2B,D). The pyrrolo pyimidine core for each compound, as expected, was predicted to form H-bonds with Asp^22^ and Ile^23^. Compound **6g**, which has the same number atoms between the amine and the carboxyl group, is predicted to take advantage of the same hydrogen bonds with Phe^156^ and Asp^157^ that were observed with compound **1** (Figure 2A). Compound **6h**, which has an extra carbon between the amine and carboxyl, is unable to make that interaction (Figure 2C). Interestingly, it is predicted to stretch the carboxylic acid into the phosphate binding groove, forming the same hydrogen bonds previously observed with the ADPr-bound structure, including one of the conserved water molecules within that groove.

**Figure 2.**
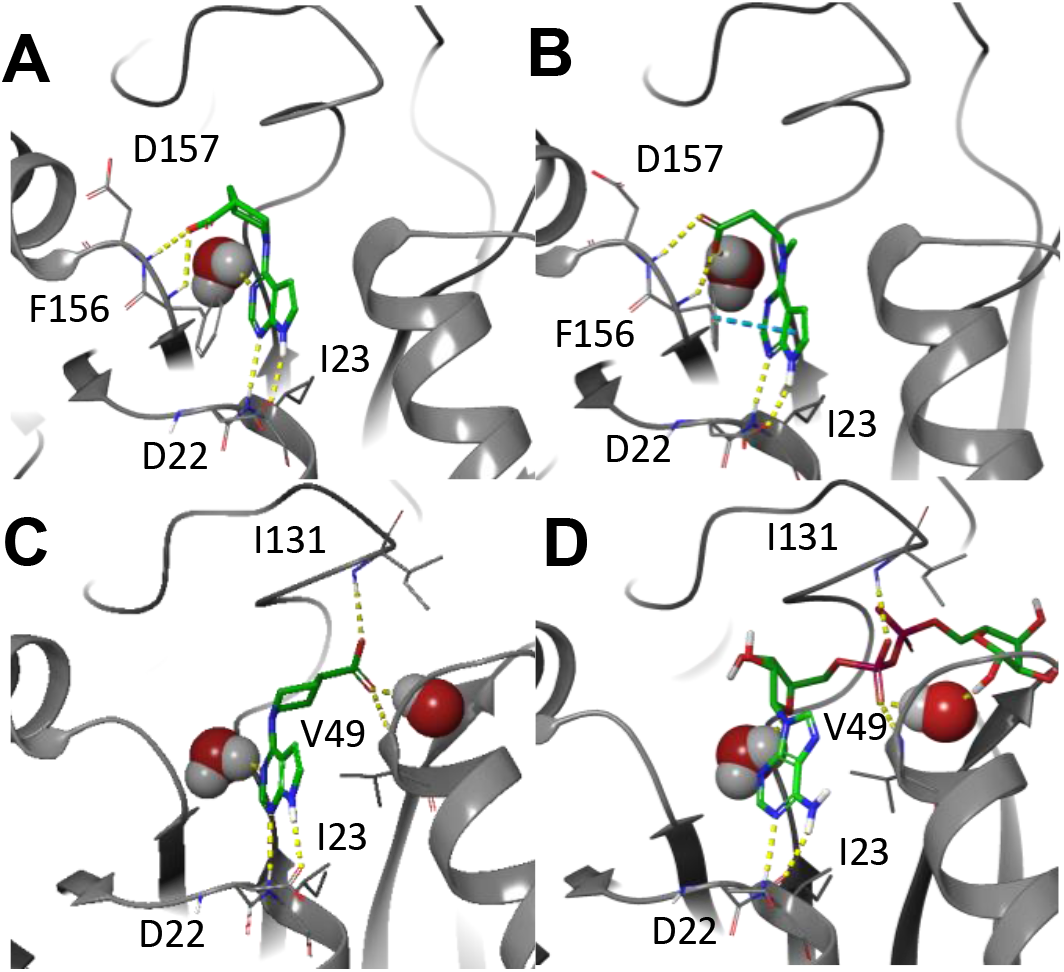
Models of compounds 6g and 6h in the Mac1 active site. A and C) Compound **6g** and **6h** (C) were docked into several Mac1 structures, including the co-crystallized structure with parent compound **1**. B and D) and 6WOJ, the co-crystallized structure with ADPr. Hydrogen bonds are represented as yellow dashes, while pi-pi interactions are represented as cyan dashes. Compound **6g** adopts a very similar binding pose to parent compound **1**, while the extra carbon between the amine and carboxyl group in compound **6h** results in different interactions for the carboxylate that strongly resemble those of the phosphate of ADPr.

Primary amino acid derivatives as synthesized in Schemes 3 and 4 were also analyzed in the AS assay for Mac1 binding (Table 2). The simplest amino acid derivative, glycinate **15d** did not demonstrate any inhibitory effect on binding in the AS assay at the highest concentration tested. Some binding activity was noted with the valine ethyl ester derivative **14a** (IC_50_ = 45 μM) and the valinate **15a** (IC_50_ = 24 μM). Interestingly, the closely related leucine derivatives **14b/15b** were completely inactive indicating that there may be a negative steric effect from the isobutyl group. The most potent derivative from the natural amino acids was the tryptophan derivative. The tryptophanate **15c** was in fact the most potent compound discovered in either series (IC_50_ = 6.1 μM in AS) (Figure 4A). We confirmed its potency with a FRET assay measuring binding of a CFP-Mac1 to cysteine MARylated GAP-TAG fused to YFP [22]. The measured IC_50_ (11.1 ± 3.19 μM) was in agreement with the AS assay (Fig. 4B). Molecular modeling was performed for **15c** is shown in Figure 3 along with the starting fragment **1**. The pyrrolo pyrimidine core of **15c** once again is predicted to form H-bonds with Asp^22^ and Ile^23^ in the Adenosine binding pocket (Figure 3A) similar to fragment **1** (Figure 3B). The carboxylate of **15c** also formed a hydrogen bond network with Phe^156^, Asp^157^ and a water molecule. The indole ring of **15c** was predicted to be positioned in the phosphate binding pocket, adding van der Waals interactions while essentially locking the carboxylate into position. Outlining the importance of the carboxylic acid, tryptamine derivative **20** was approximately 7-fold less potent than **15c**. Beta amino acid derivatives **18** and **19** also demonstrate potency comparable to the best secondary amino acid derivatives with the methyl ester **18** (IC_50_ = 30 μM) almost as potent as the corresponding carboxylic acid **19** (IC_50_ = 25.2 μM).

**Table 2.**
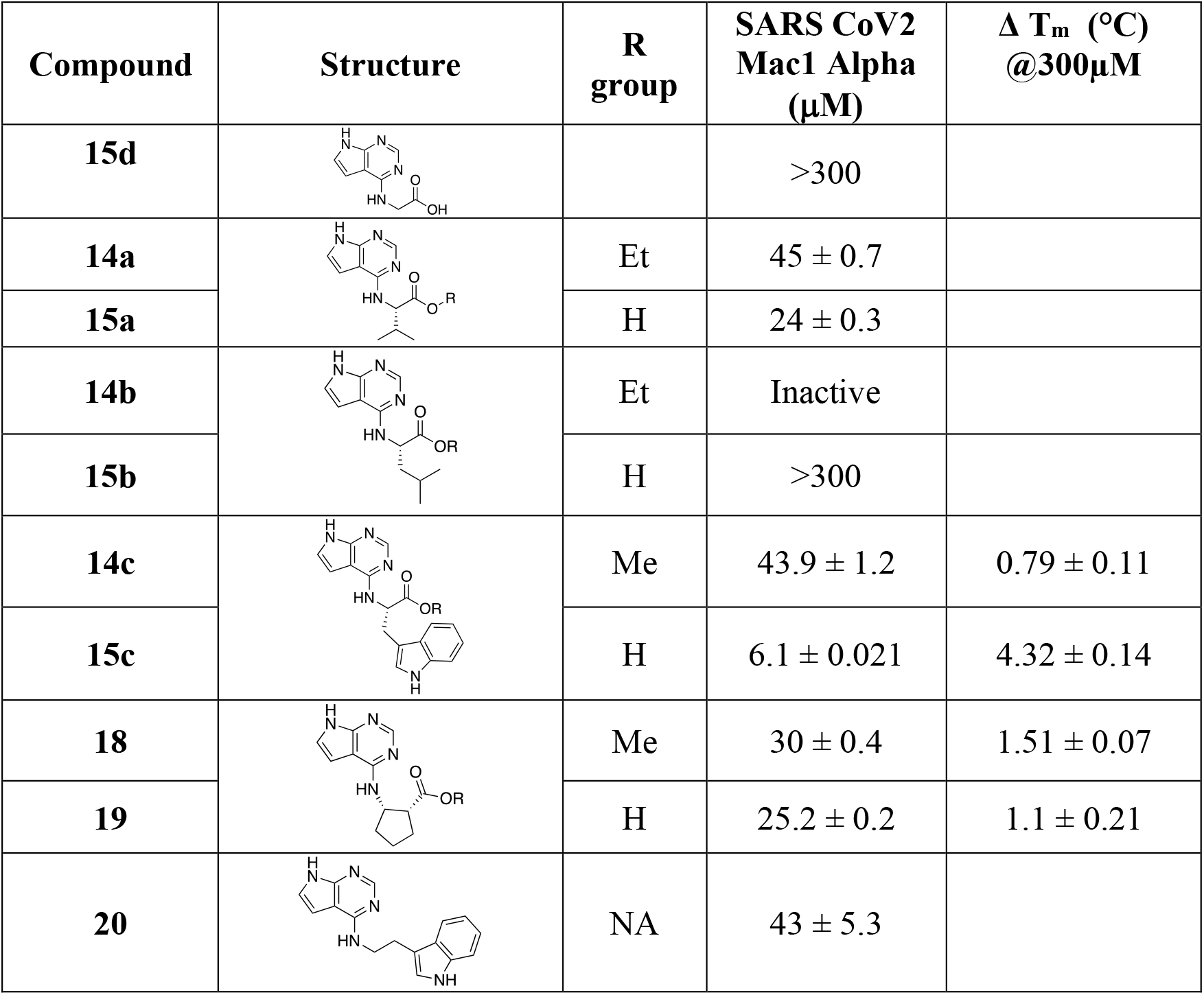
Inhibition data for primary amino acid derivatives. IC_50_s, changes in the melting temperature and corresponding SDs of three replicates are listed.

| Compound | Structure | R group | SARS CoV2 Mac1 Alpha (μM) | Δ T<sub>m</sub> (°C) @300μM |
| --- | --- | --- | --- | --- |
| **15d** |  |  | >300 |  |
| **14a** |  | Et | 45 ± 0.7 |  |
| **15a** |  | H | 24 ± 0.3 |  |
| **14b** |  | Et | Inactive |  |
| **15b** |  | H | >300 |  |
| **14c** |  | Me | 43.9 ± 1.2 | 0.79 ± 0.11 |
| **15c** |  | H | 6.1 ± 0.021 | 4.32 ± 0.14 |
| **18** |  | Me | 30 ± 0.4 | 1.51 ± 0.07 |
| **19** |  | H | 25.2 ± 0.2 | 1.1 ± 0.21 |
| **20** |  | NA | 43 ± 5.3 |  |

**Figure 3.**
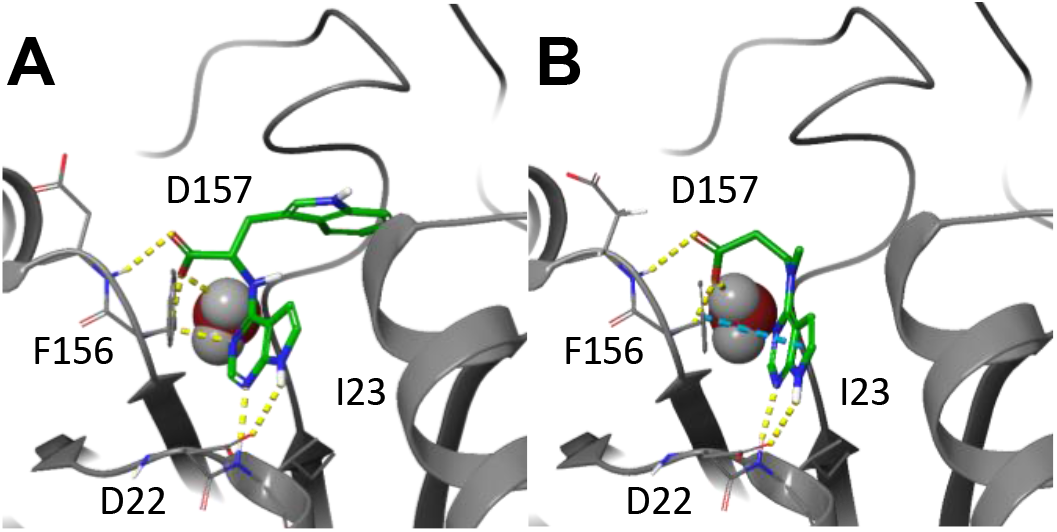
Model of compound 15c and fragment 1 in the active site of Mac1. Compound **15c** (A) was docked into the co-crystallized structure with parent compound **1 (**B). Hydrogen bonds are represented as yellow dashes, while pi-pi interactions are represented as cyan dashes. Compound **15c** adopts a very similar binding pose to parent compound **1**, while orienting the indole into the phosphate binding groove.

**Figure 4.**
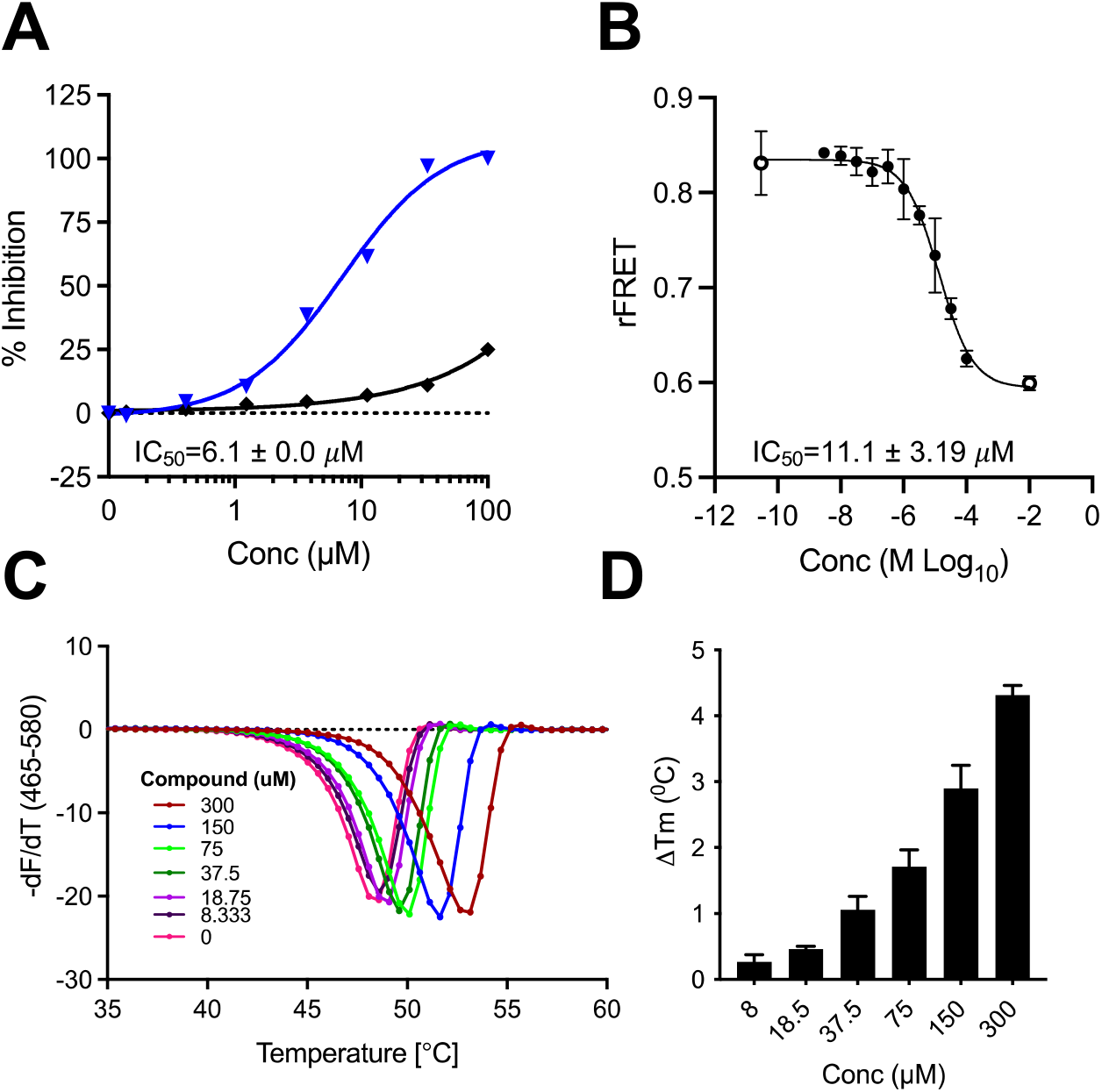
Inhibition and binding of SARS-CoV-2 Mac1-ADP-ribose binding by Compound **15c**. A) Dose response curve of Mac1-ADP-ribose binding with **15c** in the AS (n=2). B) Dose response curve of Mac1-ADP-ribose binding with **15c** in FRET assay (n=3). C-D) Stabilization of SARS-CoV-2 Mac1 by Compound **15c** in DSF assay (n=2).

Evidence for direct Mac1 binding was obtained by using differential scanning fluorimetry (DSF) thermal shift analysis of our most potent compounds from these two series. As shown in Table 3, compounds **6h** and **19** demonstrated a thermal shift of >1 °C, while the T_m_ for compound **6g** was ~1.7 °C. The most significant thermal shift was noted with our most potent compound, **15c**, and was 4.3 °C (Figure 4C-D), comparable to ADPr (T_m_ = 4.1C).

To further demonstrate the potential for this series of compounds to inhibit the enzymatic activity of Mac1, **6g** and **15c** were assayed using a recently developed ADPr-Glo assay [18]. This assay utilizes the enzyme NudF, which is a phosphatase that cleaves free ADP-ribose into AMP and diphosphate but cannot cleave ADP-ribose attached to a protein. In this assay, both Compound **6g** and **15c** demonstrated a dose-dependent inhibition of Mac1 enzyme activity (Figure 5). Compound **15c** demonstrated greater inhibition, consistent with its increased binding activity. These results demonstrate that both **6g** and **15c** can inhibit Mac1 ADP-ribosylhydrolase activity.

**Figure 5.**
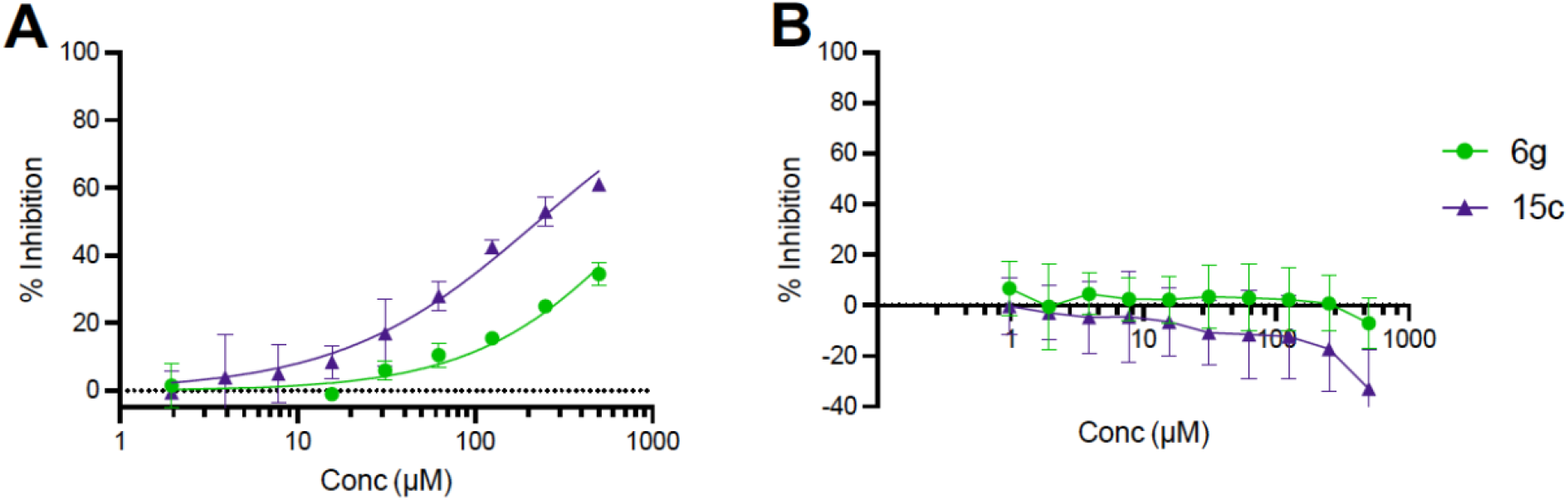
Compounds **6g** and **15c** inhibit SARS-CoV-2 Mac1-ADP-ribosylhydrolase activity. A) Dose response curve for **6g** and **15c** against SARS-CoV-2 Mac1 in the ADPr-Glo assay (n=2). B) Neither compound demonstrated any activity in a NudF-mediated counterscreen.

**Figure 6.**
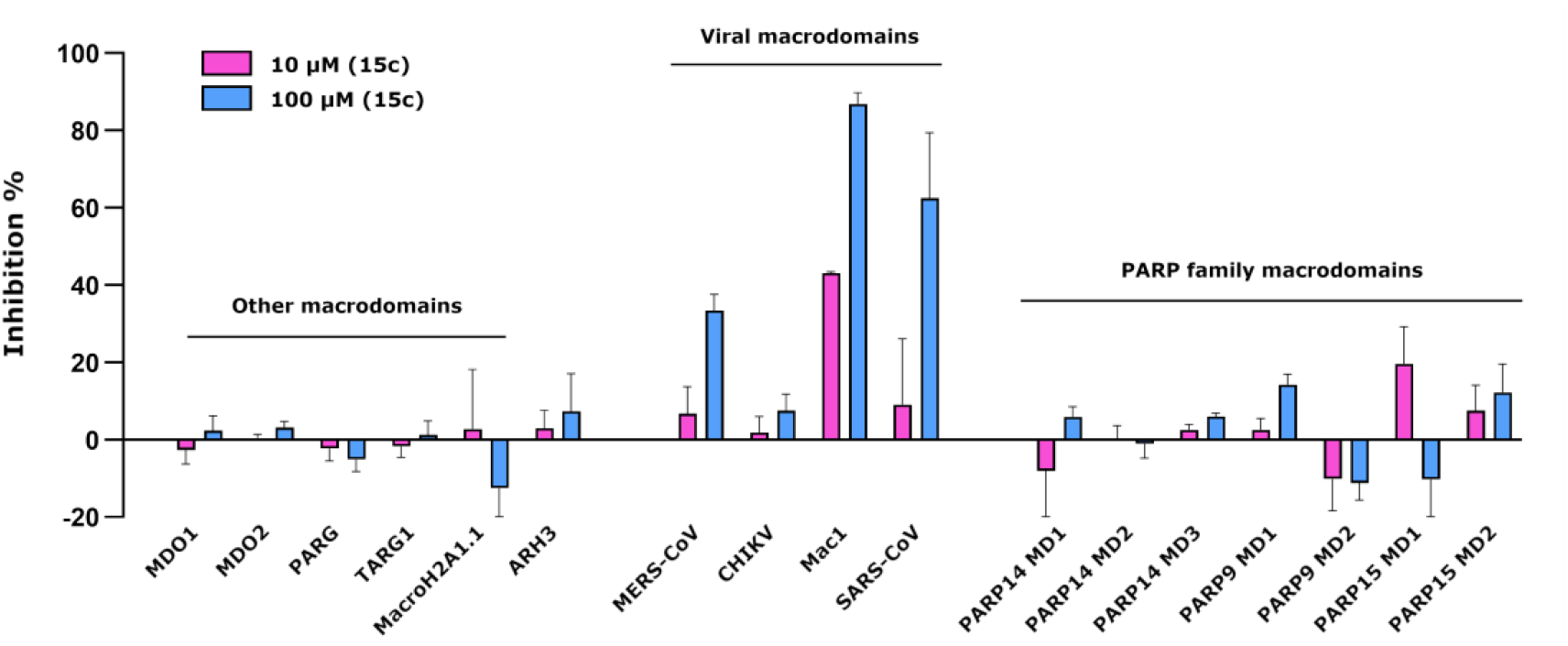
Inhibition profile for **15c** against a panel of ADP-ribose binding proteins. Data shown are means and standard deviations of four replicates.

To determine the specificity of **15c**, we utilized the FRET assay and tested **15c** for its ability to inhibit MAR binding of an extended panel of MAR binding proteins from viruses and humans, including human macrodomains, the CHIKV macrodomain, and 3 separate coronavirus macrodomains. Compound **15c** was remarkably selective for coronavirus macrodomains, specifically SARS-CoV-2, as shown in the first three columns.

In conclusion, we present the design, synthesis, binding evaluation and inhibition of Mac1 by a series of primary and secondary amino acid derivatives of *7H*-pyrrolo[2,3-*d*]pyrimidines in this manuscript. Structure activity studies were conducted against the SARS-CoV-2 macrodomain (Mac1), a potential anti-viral drug target. A molecular modeling study defined a predictive binding mode for this series outlining the importance of the pyrrolo pyrimidine core and a carboxylic acid 3-4 atoms from the 4 position of the core. Low micromolar derivatives **6g** and **6h** were discovered from the secondary amino acid series that also demonstrated selectivity over human macrodomain. Compound **15c** from the primary amino acid series demonstrated low micromolar binding potency against Mac1 and a thermal shift comparable to ADPr. In addition, compounds **6g** and **15c** demonstrated the ability to inhibit the enzymatic activity of Mac1. Compound **15c** also showed a clear selectivity profile towards coronavirus macrodomains, especially towards SARS-CoV-2 Mac1, which may be important when developing the compound further to study its effects on the virus infection. Taken together, many of these derivatives represent some of the most potent small molecule inhibitors of the SARS-CoV-2 macrodomain in the literature, many of which have MW<300 leaving room for derivatization and optimization while maintaining drug like parameters.

## Abbreviations

ADPr: Adenosine diphosphate ribose
CoV: Coronavirus
SARS-CoV-2: -Severe acute respiratory syndrome coronavirus 2
COVID19: Coronavirus disease 2019
ARTs: Adenosine diphosphate ribosyl transferases
Mac1: Macrodomain 1 of SARS-CoV-2
ARH: Adenosine diphosphate ribosyl hydrolase
PAR: poly-Adenosine diphosphate ribose
PARG: poly-Adenosine diphosphate ribose glycohydrolase
MHV-JHM: Mouse hepatitis strain JHM
FRET: Fluorescence resonance energy transfer
DSF: Differential Scanning Fluorimetry
CFP: Cyan Fluorescent Protein
SAR: Structure activity relationships
YFP: Yellow Fluorescent Protein

## Acknowledgements

DVF would like to acknowledge McDaniel College Student-Faculty Summer Research Fund, the Jean Richards Fund, the Schofield fund, and the Scott and Natalie Dahne fund. DVF would also like to acknowledge Mr. Kristopher Mason and Vaccitech for the use of their mass spectrometer. LL would like to acknowledge the use of the facilities of the Biocenter Oulu Structural Biology core facility, a member of Biocenter Finland, Instruct-ERIC Centre Finland and FINStruct. This research was funded by National Institutes of Health (NIH) grants P20 GM113117, P30GM110761, a CTSA grant from NCATS awarded to the University of Kansas for Frontiers: University of Kansas Clinical and Translational Science Institute UL1TR002366 and University of Kansas start-up funds to A.R.F, and by Sidrid Jusélius foundation grant to L.L., and Johns Hopkins Bloomberg School of Public Health Discretionary Fund to A. K. L. L.

## Supporting Information

### Plasmids

SARS-CoV-2 Mac1 (residues 205-379 of nsp3) was cloned into the pET30a+ expression vector with an N-terminal His tag and a TEV cleavage site (Synbio) [24]. The pETM-CN Mdo2 Mac1 (residues 7-243) expression vector with an N-terminal His-TEV-V5 tag and the pGEX4T-PARP10-CD (residues 818-1025) expression vector with an N-terminal GST tag were previously described. All plasmids were confirmed by restriction digest, PCR, and direct sequencing.

### Protein Expression and Purification

The proteins used were prepared as previously described [22, 24, 25].

### Differential Scanning Fluorimetry (DSF)

Thermal shift assay with DSF involved use of LightCycler^®^ 480 Instrument (Roche Diagnostics). In total, a 15 μL mixture containing 8X SYPRO Orange (Invitrogen), and 10 μM macrodomain protein in buffer containing 20 mM HEPES, NaOH, pH 7.5 and various concentrations of ADP-ribose or hit compounds were mixed on ice in 384-well PCR plate (Roche). Fluorescent signals were measured from 25 to 85°C in 0.2 °C/30/Sec steps (excitation, 470-505 nm; detection, 540-700 nm). The main measurements were carried out in triplicate. Data evaluation and T_m_ determination involved use of the Roche LightCycler® 480 Protein Melting Analysis software, and data fitting calculations involved the use of single site binding curve analysis on GraphPad Prism. The thermal shift ΔTm was calculated by subtracting the Tm values of the DMSO from the Tm values of compounds.

### AlphaScreen™ Assay

The AlphaScreen™ reactions were carried out in 384-well plates (Alphaplate, PerkinElmer, Waltham, MA) in a total volume of 40 μL in buffer containing 25 mM HEPES (pH 7.4), 100 mM NaCl, 0.5 mM TCEP, 0.1% BSA, and 0.05% CHAPS. All reagents were prepared as 4X stocks and 10 μL volume of each reagent was added to a final volume of 40 μL. All compounds were transferred acoustically using ECHO 555 (Beckman Inc) and preincubated after mixing with purified His-tagged macrodomain protein (250 nM) for 30 min at RT, followed by addition of a 10 amino acid biotinylated and ADP-ribosylated peptide [ARTK(Bio)QTARK(Aoa-RADP)S] (Cambridge peptides) (625 nM). After 1h incubation at RT, streptavidin-coated donor beads (7.5 μg/ml) and nickel chelate acceptor beads (7.5 μg/mL); (PerkinElmer AlphaScreen™ Histidine Detection Kit) were added under low light conditions, and plates were shaken at 400 rpm for 60 min at RT protected from light. Plates were kept covered and protected from light at all steps and read on BioTek plate reader using an AlphaScreen™ 680 excitation/570 emission filter set. For counter screening of the compounds, 25 nM biotinylated and hexahistidine-tagged linker peptide (BIO-6His) (PerkinElmer) was added to the compounds, followed by addition of beads as described above.

### ADPr Glo Assay

A recently developed luminescence based ADP-ribosylhydrolase assay was used to measure the ability of specific compounds to inhibit Mac1 ADP-ribosylhydrolase activity. Briefly, MARylated PARP10-CD was incubated with the SARS-CoV-2 Mac1 (0.86 nM) and NudF (125 nM) proteins at room temperature for 30 min. Proteins were removed with a Microcon spin filter (10,000 MWCO) and the products were measured with AMP-Glo. Reactions without Mac1 were performed in parallel as a negative control. Also, NudF was incubated with free ADP-ribose as a counter screen. Luminescence signal was converted to AMP concentration via interpolation from a standard curve. Data plotted are AMP generated by the macrodomain and NudF, subtracted by AMP generated from NudF alone. Inhibition percentages were calculated and non-linear regression analysis was performed in GraphPad Prism.

### FRET assay

A FRET method was utilized for the profiling of **15c** a panel of human and viral macrodomains to determine their specificity [22, 25, 26]. The assay is based on the site-specific introduction of cysteine-linked mono-ADP ribose to the C-terminal Gαi peptide (GAP) by Pertussis toxin subunit1 (PtxS1) fused to YFP. To generate the FRET signal ADP-ribosyl binders were fused to CFP. Samples were prepared in the assay buffer (for most binders; 10 mM Bis-Tris propane pH 7.0, 3 % (w/v) PEG 20,000, 0.01 % (v/v) Triton X-100 and 0.5 mM TCEP), (for TARG1; 10 mM Bis-Tris propane pH 7.0, 150 mM NaCl, 0.01 % (v/v) Triton X-100 and 0.5 mM TCEP), (for PARG and PARP15 MD1; 10 mM Bis-Tris propane pH 7.0, 25 mM NaCl, 0.01 % (v/v) Triton X-100 and 0.5 mM TCEP) in a 384-well black polypropylene flat-bottom plates (Greiner, Bio-one) with 10 μl reaction volume per well. The reactions consisted of 1 μM CFP-fused binders and 5 μM MARylated YFP-GAP. Reactions were excited at 410 nm (20 nm bandwidth), while the emission signal was measured at 477 nm (10 nm bandwidth) and 527 nm (10 nm bandwidth). Afterwards, blank was deducted from the individual values and the radiometric FRET (rFRET) was calculated by dividing the fluorescence intensities at 527 nm by 477 nm. Compound was dispensed with Echo acoustic liquid dispenser (Labcyte, Sunnyvate, CA). Dispensing of larger volumes of the solutions was carried out by using Microfluidic Liquid Handler (MANTIS®, Formulatrix, Beford, MA, USA). Measurements were taken with Tecan Infinite M1000 pro plate reader.

### Modeling details

Compounds were docked into the ADPr-bound (6WOJ), 3 unique unbound conformations (7KR0, 7KR1, 6WEY) and two small molecule bound (5RSG, 5RTT) structures of Mac1 [20, 24, 27]. The proteins and ligands were prepared using Schrodinger Maestro and were subsequently docked using Glide with XP precision, followed by a Prime MM-GBSA minimization, allowing flexibility for any residue within 5 Å of the ligand [28–33].

### Reagents and Methods for Chemical Purification

All solvents were reagent grade or high performance liquid chromatography (HPLC) grade. Unless otherwise noted, all materials were obtained from commercial suppliers and used without further purification. ^1^H NMR spectra were recorded at 400.19 MHz. All ^13^C spectra were recorded at 100.63 MHz. The HPLC solvent system consisted of distilled water and acetonitrile, both containing 0.1% formic acid. Analytical liquid chromatography / mass spectrometry (LC-MS) (LC/MS) were performed utilizing an Agilent Infinity 1260 DAD G7115A detector coupled with a 6120 quadrupole analyzer. Samples were run on a 5-95% gradient ACN in H_2_O over 6 minutes (both mobile phases modified w/ 0.1% TFA). All results depicted herein are (M+H)^+^ as ionization source is ESI. All final compounds tested were confirmed to be of ≥95% purity by the HPLC methods described above.

#### Methods for Chemical Synthesis

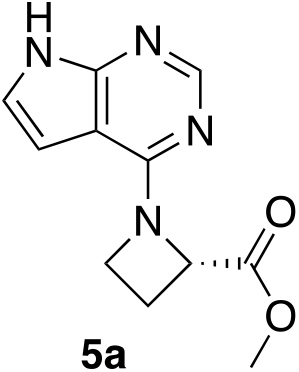

#### Methyl (*S*)-1-(*7H*-pyrrolo[2,3-*d*]pyrimidin-4-yl)azetidine-2-carboxylate (5a)

4-Chloro-*7H*-pyrrolo[2,3-*d*]-pyrimidine **3** (0.500g, 3.26 mmol, (*S*)-Azetidine 2-carboxilic acid methyl ester HCl **4a** (0.600g, 3.91mmol), isopropanol (4 mL), and DIEA (1.71mL, 9.81mmol) were combined in a 25mL round bottom and left to stir under heat for 9 days. TLC showed that the reaction was complete, therefore the isopropanol was removed via rotary evaporator. EtOAc (10mL) was added to the reaction mixture, and poured into a separatory funnel. It was washed twice with 10% aqueous sodium bicarbonate and the organic layer was dried with MgSO_4_.The product was purified through a column chromatography on silica gel and was characterized as the desired product. Dry yield: 0.175g (23%). ^1^HNMR (CDCl_3_): 11.6 (br s, 1H), 8.35 (s, 1H), 7.09 (d, 1H, *J* = 3.6Hz), 6.34 (d, 1H, *J* = 3.6Hz), 5.11 (m, 1H), 4.54 (m, 1H), 4.38 (m, 1H), 3.79 (s, 3H), 2.82 (m, 1H), 2.50 (m, 1H); ^13^C NMR (CDCl_3_): 172.51, 156.67, 151.46, 151.35, 121.68, 102.68, 99.45, 61.86, 52.62, 49.99, 22.43; MS (ES+, M+1): 233.2.

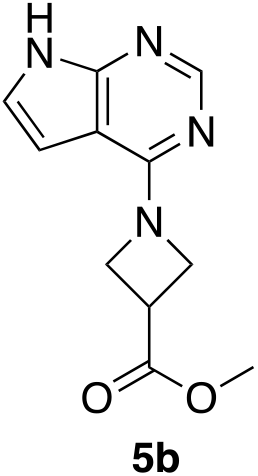

#### Methyl 1-(*7H*-pyrrolo[2,3-*d*]pyrimidin-4-yl)azetidine-3-carboxylate (5b)

1-4-Chloro-7*H*-pyrrolo[2,3-*d*]-pyrimidine **3** (0.300g, 1.961 mmol), methyl-azetidine-3-carboxylate-hydrochloride **4b** (0.356g, 2.35mmol), isopropanol (3mL) and DIEA (1.02mL, 5.88mmol) were combined in a 25mL round bottom flask and stirred at 70 °C for 24h. TLC indicated that reaction was complete. The reaction was concentrated in vacuo and the reaction mixture was diluted with ethyl acetate (10mL) and washed with 1M HCl (3mL) and 10% NaHCO_3_ (2 x 3 mL). The organic layer was dried with MgSO_4_, filtered and chromatographed (EtOAc/Hexanes) to afford the desired product as a white solid with dry yield of 0.237g, 52.1%. ^1^HNMR (CDCI_3_): 11.20 (s, 1H), 8.33 (s, 1H), 7.09 (d, 1H, *J* = 3.2Hz), 6.39 (d, 1H, *J* = 3.2Hz), 4.57 (d, 4H, *J* = 8.8Hz), 3.78 (s, 3H), 3.67 (m, 1H); ^13^C NMR (DMSO-*d_6_*): 172.93, 156.77, 150.95, 150.67, 121.95, 101.75, 98.83, 53.21, 52.02, 33.14; MS (ES+, M+1): 233.2.

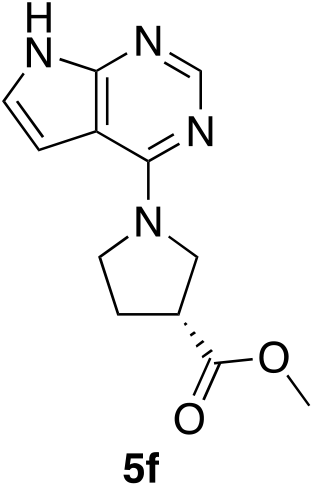

#### Methyl (*R*)-1-(*7H*-pyrrolo[2,3-*d*]pyrimidin-4-yl)pyrrolidne-3-carboxylate (5f)

A 25.0 mL round bottom was used to combine chloride **3** (0.501 g, 3.27 mmol), (*R*)-methyl-pyrrolidine-3-carboxylate, HCl **4f** (0.506 g, 3.92 mmol), DIEA (1.71 mL, or 9.81 mmol), and 3 mL of isopropanol. The reaction was heated for 24 hours, a TLC indicated that the reaction was complete (EtOAc). The reaction was concentrated on the rotary evaporator to remove the isopropyl alcohol. EtOAc (10 mL) was added to the round bottom. The organic layer was washed with 10% NaHCO_3_ and the organic layer was dried with MgSO_4_, filtered off using vacuum filtration, chromatographed (EtOAc/hexanes). Dry yield = 0.475g (59%). ^1^HNMR (CDCl_3_): δ 10.91 (s, 1H), 8.33 (s, 1H), 7.08 (d, 1H, *J* = 4.0Hz), 6.59 (d, 1H, *J* = 4.0Hz), 4.13 (m, 4H), 3.89 (s, 3H), 3.25 (m, 1H), 2.35 (m, 2H); ^13^C NMR (DMSO-*d_6_*): 173.46, 154.59, 151.09, 150.96, 120.75, 102.42, 100.60, 51.85, 49.48, 46.75, 41.79, 28.08; MS (ES+, M+1): 247.1.

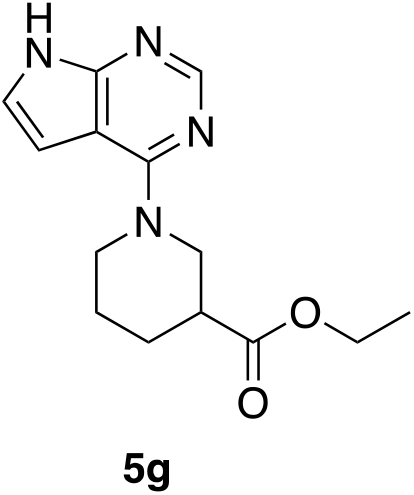

#### Ethyl 1-(*7H*-pyrrolo[2,3-*d*]pyrimidin-4-yl)piperidine-3-carboxylate (5g)

In a 25 mL round bottom flask 500 mg of the chloride **3** (3.26 mmol), 0.56 mL of racemic ethyl-piperidine-3-carboxylate HCl **4g** (3.6 mmol), and 890 mg of K_2_CO_3_ (6.53 mmol) were dissolved in 2 mL of DMF and heated to ~140°C for 1 hour. A small sample of the reaction was diluted in DCM and a TLC of the reaction mixture (EtOAc) indicated that the starting material was consumed and a lower running spot was apparent. The mixture was cooled to rt for 5 min. Water (6 mL) was added to the round bottom and the mixture was stirred. A tan solid formed almost immediately and the reaction continued to stir for 10 min. The solid was filtered off and washed with dH_2_O (~10-20mL). The solid was air dried under vacuum for another 5 minutes and then transferred to a vial to dry overnight. Dry yield = 0.65g (72.9%). ^1^HNMR (DMSO-*d_6_*): δ 11.69 (s, 1H), 8.13 (s, 1H), 7.18 (dd, 1H, *J* = 3.6, 2.0Hz), 6.55 (dd, 1H, *J* = 3.6, 2.0Hz), 4.62 (dd, 1H, *J* = 13.6, 3.6Hz), 4.31 (d, 1H, *J* = 13.6Hz), 4.06 (q, 2H, *J* = 6.8Hz), 3.34 (m, 2H), 2.54 (m, 1H), 1.97 (m, 1H), 1.76 (m, 2H), 1.52 (m, 1H), 1.15 (t, 3H, *J* = 6.8Hz). ^13^C NMR (DMSO-*d_6_*): 172.83, 156.22, 152.01, 150.60, 121.45, 102.19, 100.73, 60.06, 47.19, 45.81, 40.51, 26.79, 23.79, 14.04; MS (ES+, M+1): 275.1.

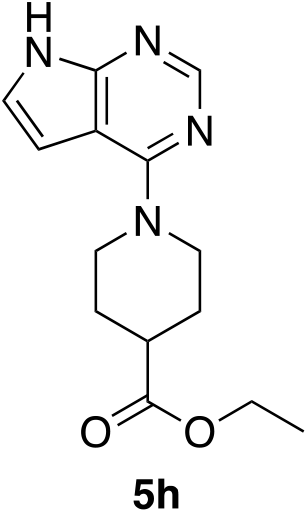

#### Ethyl 1-(*7H*-pyrrolo[2,3-*d*]pyrimidin-4-yl)piperidine-4-carboxylate (5h)

In a 25 mL round bottom flask 500 mg of the chloride **3** (3.26 mmol), 0.56 mL of ethyl piperidine-4-carboxylate **4h** (3.6 mmol), and 675 mg of K_2_CO_3_ (4.89 mmol) were dissolved in 2 mL of DMF and heated to ~140°C for 3 hours. A small sample of the reaction was diluted in DCM and a TLC of the reaction mixture (EtOAc) indicated that the starting material was consumed and a lower running spot was apparent. The mixture was cooled to rt for 5 min. Water (6 mL) was added to the round bottom and the mixture was stirred. A tan solid formed almost immediately and the reaction continued to stir for 10 min. The solid was filtered off and washed with dH_2_O (~10-20mL). The solid was air dried under vacuum for another 5 minutes and then transferred to a vial to dry overnight. Dry yield = 0.657g (72.6%). ^1^HNMR (CDCl_3_): δ 10.82 (bs, 1H), 8.34 (s, 1H), 7.11 (d, 1H, *J* = 3.6Hz), 6.52 (d, 1H, *J* = 3.6), 4.67 (d, 2H, *J* = 13.0Hz), 4.15 (q, 2H, *J* = 7.2Hz), 3.30 (t, 2H, *J* = 13.0Hz), 2.65 (m, 1H), 2.05 (m, 2H), 1.85 (m, 2H), 1.26 (t, 3H, *J* = 7.2Hz); ^13^C NMR (DMSO-*d_6_*): 173.97, 156.22, 151.94, 150.57, 121.31, 102.17, 100.77, 59.87, 44.57, 40.25, 27.63, 14.03; MS (ES+, M+1): 275.1.

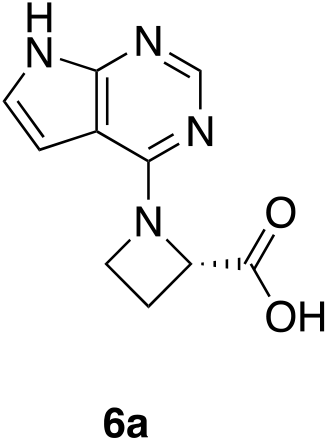

#### (*S*)-1-(*7H*-pyrrolo[2,3-*d*]pyrimidin-4-yl)azetidine-2-carboxylic acid (6a)

Methyl (*S)-1-(7H-* pyrrolo[2,3-*d*]pyrimidin-4-yl)azetidine-2-carboxylate **5a** (152 mg, 0.534 mmol), 3.0 M HCl (3 mL) and a stir bar were placed in a 25 mL RBF and heated to reflux for two hours. The solution was concentrated in vacuo until a brown solid could be observed in the flask. The residue was allowed to dry overnight at RT and the following morning the residue was placed in a vacuum oven at 110 F for 2 hours, triturated with 5 mL of Et_2_O, vacuum filtered and allowed to sit undisturbed in a vacuum oven at room temperature for another day. Dry yield 74 mg (54% yield). ^1^HNMR (DMSO-*d_6_*): 12.84 (s, 1H), 8.35 (m, 1H), 7.48 (br s, 1H), 6.7 (br s, 1H), 5.31 (m, 1H), 4.46 (m, 2H), 2.89 (m, 1H), 2.46 (m, 1H); ^13^C NMR (DMSO-*d_6_*): 171.55, 149.45, 145.36, 142.39, 125.21, 101.82, 100.41, 63.50, 51.46, 21.62; MS (ES+): MS (ES+, M+1): 219.1.

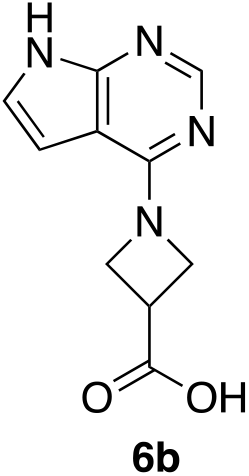

#### 1-(*7H*-pyrrolo[2,3-*d*]pyrimidin-4-yl)azetidine-3-carboxylic acid (6b)

Ester **5b** (0.20g, 0.86 mmol) was suspended in 3M HCl (3mL) in a 25mL round bottom flask and heated to reflux for 30min. TLC indicated that reaction was complete. Removal of solvent and trituration with a minimal amount of cold water afforded the desired acid. Yield = 131mg (70%). ^1^HNMR (DMSO-*d_6_*): δ 12.88 (s, 1H), 8.28 (s, 1H), 7.45 (dd, 1H, *J* = 3.6, 2.0Hz), 6.78 (dd, 1H, *J* = 3.6, 2.0Hz), 4.4-4.8 (m, 4H), 3.72 (m, 1H); ^13^C NMR (DMSO-*d_6_*): 172.85, 149.63, 146.43, 142.55, 124.83, 101.87, 100.35, 55.33, 32.97; MS (ES+, M+1): 219.1.

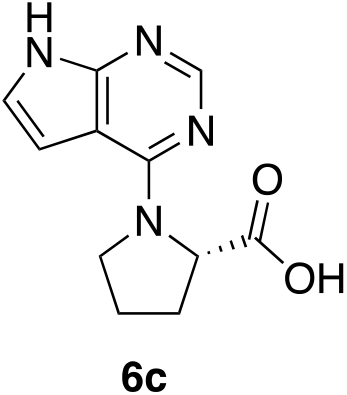

#### (*7H*-pyrrolo[2,3-*d*]pyrimidin-4-yl)-*Z*-proline (6c)

A 25.0 mL round bottom was used to combine the chloride **3** (0.499 g, 3.26 mmol), *L*-proline (0.451 g, 3.91 mmol), DIEA (1.70 mL, 9.78 mmol), and 3.0 mL of isopropanol. The reaction was heated to ~70 °C for 48 hours, TLC indicated that the reaction was complete (EtOAc). The reaction was concentrated on the rotary evaporator to remove the isopropyl alcohol. The product was partitioned between 5 mL of EtOAc and 3 mL 10% NaHCO_3_. The bicarbonate layer was acidified to a pH of 3 with 3M HCl, then extracted again with boiling EtOAc. Product began to crystallize in the aqueous layer, so vacuum filtration was then used to retrieve the final product. After drying, it was characterized by dry yield = 0.154 g, 20.3%. ^1^HNMR (DMSO-*d_6_*): δ 12.50 (bs, 1H), 11.61 (s, 1H), 8.04 (s, 1H), 7.13 (dd, 1H, *J* = 3.6, 2.0Hz), 6.56 (br s, 1H), 4.64 (m, 1H), 3.91 (m, 2H), 2.01-2.24 (m, 4H); ^13^C NMR (DMSO-*d_6_*): 174.36, 154.42, 151.11, 150.82, 121.09, 102.51, 100.62, 59.85, 48.18, 28.77, 24.31; MS (ES+, M+1): 233.1.

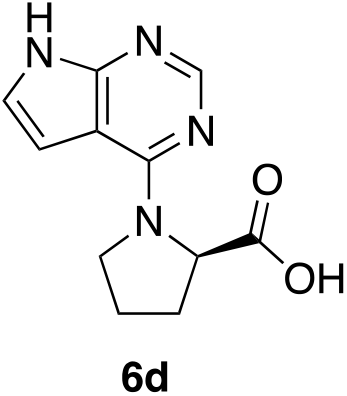

#### (*7H*-pyrrolo[2,3-*d*]pyrimidin-4-yl)-*D*-proline (6d)

This compound was made in an identical manner to **6c** using *D*-Proline as the starting material. Dry yield = 0.040 g, 29.2%. ^1^HNMR (DMSO-*d_6_*): δ 12.50 (bs, 1H), 11.61 (s, 1H), 8.04 (s, 1H), 7.13 (dd, 1H, *J* = 3.6, 2.0Hz), 6.56 (br s, 1H), 4.64 (m, 1H), 3.91 (m, 2H), 2.01-2.24 (m, 4H); ^13^C NMR (DMSO-*d_6_*): 174.36, 154.42, 151.11, 150.82, 121.09, 102.51, 100.62, 59.85, 48.18, 28.77, 24.31; MS (ES+, M+1): 233.1.

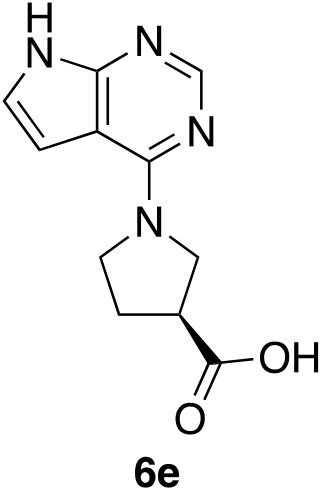

#### (*S*)-1-(*7H*-pyrrolo[2,3-*d*]pyrimidin-4-yl)pyrrolidine-3-carboxylate (6e)

This reaction was conducted in a similar manner to **6c** using (S) pyrrolidine-3-carboxylic acid as the starting material. Dry weight = 424mg (56%). ^1^HNMR (DMSO-*d_6_*): δ 11.57 (bs, 1H), 8.06 (s, 1H), 7.08 (dd, 1H, *J* = 3.6, 2.0Hz), 6.53 (dd, 1H, *J* = 3.6, 2.0Hz), 3.80 (m, 4H), 3.15 (m, 1H), 2.18 (m, 2H); ^13^C NMR (DMSO-*d_6_*): 173.55, 148.46, 147.01, 142.68, 124.09, 103.92, 101.41, 51.39, 49.30, 42.72, 28.38; MS (ES+, M+1): 233.1.

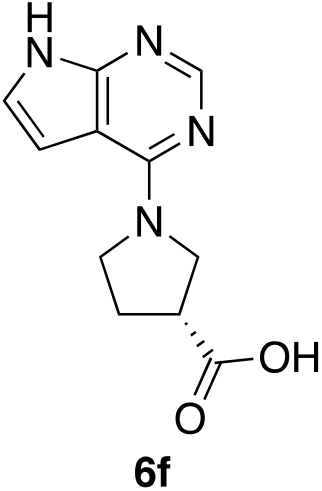

#### (*R*)-1-(*7H*-pyrrolo[2,3-*d*]pyrimidin-4-yl)pyrrolidine-3-carboxylic acid (6f)

A 25.0 mL round bottom was used to combine the ester **5f** (0.464 g, 1.88 mmol) and 5 mL of 3 M HCl. The reaction was heated to reflux for 2 hours. It was then cooled and a white precipitate crashed out of solution and was filtered using vacuum filtration. Dry weight = 0.322 g, 69.4%. ^1^HNMR (DMSO-*d*): δ 11.6 (br s, 1H), 8.06 (s, 1H), 7.08 (dd, 1H, *J* = 3.6, 2.0Hz), 6.53 (dd, 1H, *J* = 3.6, 2.0Hz), 3.80 (m, 4H), 3.15 (m, 1H), 2.18 (m, 2H); ^13^CNMR (DMSO-*d_6_*): 173.55, 148.46, 147.01, 142.68, 124.09, 103.92, 101.41, 51.39, 49.30, 42-72, 28.38; MS (ES+, M+1): 233.1.

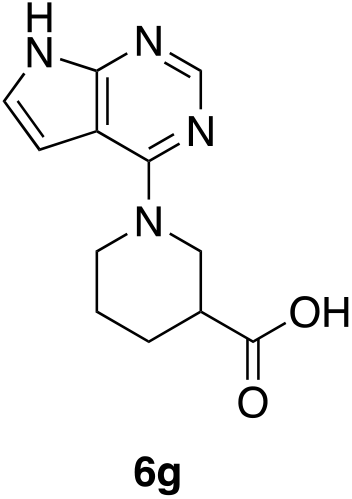

#### 1-(*7H*-pyrrolo[2,3-*d*]pyrimidin-4-yl)piperidine-3-carboxylic acid (6g)

In a 25 mL round bottom flask, the ethyl ester **5g** (230 mg, 0.84 mmol) was dissolved in a mixture of methanol (2 mL) and 5% NaOH (2 mL). This mixture was heated to ~70 °C for 1 hour. The reaction was cooled to rt then the methanol was removed via rotary evaporation leaving the product in ~2 mL of aqueous base. To this mixture was added 1 mL of water and 5 mL of EtOAc. The bilayer was separated using a separatory funnel and the bottom aqueous layer was acidified to pH 3 using 3M HCl. The acidic aqueous layer was extracted with EtOAc (3 x 10 mL). The combined organics were dried and concentrated to afford a white crystalline solid that was triturated with Et_2_O and collected by vacuum filtration. Dry yield = 0.11g (52%) ^1^HNMR(DMSO-*d_6_*): δ 12.39 (s, 1H), 11.69 (s, 1H), 8.13 (s, 1H), 7.17 (dd, 1H, *J* = 3.6, 2.0Hz), 6.54 (dd, 1H, *J* = 3.6, 2.0Hz), 4.66 (dd, 1H, *J* = 13.2, 4.0Hz), 4.40 (d, 1H, *J* = 13.2Hz), 3.22 (m, 2H), 2.46 (m, 1H), 1.97 (m, 1H), 1.71 (m, 2H), 1.49 (m, 1H); ^13^C NMR (DMSO-*d_6_*): 174.58, 156.31, 151.99, 150.63, 121.43, 102.24, 100.73, 47.51, 45.82, 40.68, 26.95, 24.05; MS (ES+, M+1): 247.2.

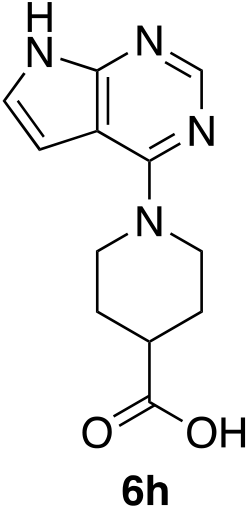

#### 1-(*7H*-pyrrolo[2,3-*d*]pyrimidin-4-yl)piperidine-4-carboxylic acid (6h)

In a 25 mL round bottom flask, the ethyl ester **5h** (320 mg, 1.17 mmol) was dissolved in a mixture of methanol (2 mL) and 5% NaOH (2 mL). This mixture was heated to ~70 °C for 1 hour. The reaction was cooled to rt then the methanol was removed via rotary evaporation leaving the product in ~2 mL of aqueous base. To this mixture was added 1 mL of water and 5 mL of EtOAc. The bilayer was separated using a separatory funnel and the bottom aqueous layer was acidified to pH 3 using 3M HCl. Crystals precipitated out of solution and the mixture was cooled in an ice bath for 5 minutes then filtered and washed with 1 mL of cold water. The solid was dried overnight to afford 0.19g, 65%. ^1^HNMR (DV1SO-*d_6_*): δ 12.25 (s, 1H), 11.67 (s, 1H), 8.11 (s, 1H), 7.15 (dd, 1H, *J* = 3.6, 2.4Hz), 6.56 (d, 1H, *J* = 3.6, 2.4Hz), 4.52 (d, 2H, *J* = 13.2Hz), 3.20 (dt, 2H, *J* = 12.4, 2.4Hz), 2.56 (m, 1H), 1.91 (dd, 2H, *J* = 13.2, 3.6Hz), 1.52 (m, 2H); ^13^CNMR (DMSO-*d_6_*): 175.72, 156.25, 151.93, 150.58, 121.24, 102.14, 100.81, 44.70, 40.32, 27.74; MS (ES+, M+1): 247.2.

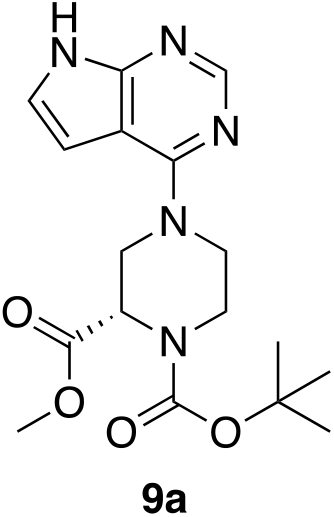

#### 1-(*tert*-butyl) 2-methyl (*S*)-4-(*7H*-pyrrolo|2,3-*d*|pyriinidin-4-yl)piperazine-1,2-dicarboxylate (9a)

4-chloro-*7H*-pyrrolo[2,3-*d*]pyimidine **3** (1.0g, 6.53mmol), DIEA (3.42mL, 19.61mmol), i-PrOH (6 mL) and (*S*)-1-*N*-boc-piperazine-2-carboxylic acid methyl ester **8a** (2.4g, 9.8 mmol) were combined into a 100 mL round bottom flask and heated to reflux for 48h. TLC (EtOAc) indicated that the reaction was complete. The i-PrOH was removed by rotary evaporation. The remaining reaction mixture was diluted in 15mL of EtOAc, poured into a 125 mL separatory funnel and washed with 10% NaHCO_3_. It was dried with anhydrous MgSO_4_, filtered and chromatographed (70/30 EtOAc/Hexanes to 100% EtOAc) to isolate the product. Dry yield = 1.72g (73%). ^1^HNMR (CDCl_3_): δ 11.05 (br s, 1H), 8.36 (s, 1H), 7.15 (d, 1H, *J* = 4.0Hz), 6.63 (br s, 1H), 5.10 (dd, 1H, *J* = 51.2, 13.6Hz), 4.91 (s, 0.5H), 4.71 (s, 0.5H), 4.61 (m, 1H), 3.98 (dd, 1H, *J* = 30.2, 12.4Hz), 3.67 (s, 3H), 3.20-3.51 (m, 4H); ^13^C NMR (DMSO-*d_6_*): {rotamer signals} {171.11, 170.77}, 156.28, {154.78, 154.26}, 151.91, 150.34 {121.91, 121.81}, 102.41, {100.49, 100.37}, {79.88, 79.67}, {55.18, 53.80}, 52.04, {45.54, 44.93}, 44.52, 41.14, {27.93, 27.80}; MS (ES+): MS (ES+, M+1): 362.2.

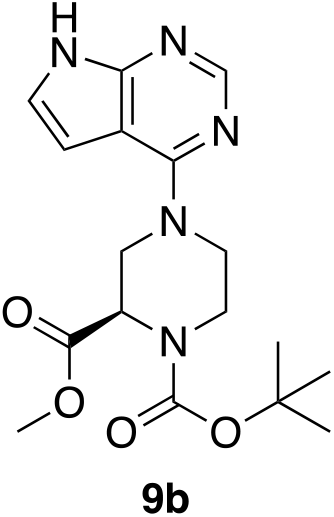

#### 1-(*tert*-butyl) 2-methyl (*R*)-4-(*7H*-pyrrolo[2,3-*d*]pyrimidin-4-yl)piperazine-1,2-dicarboxylate (9b)

This compound was made in an identical manner to **9a** using (*R*)-1-*N*-boc-piperazine-2-carboxylic acid methyl ester **8b**. Dry yield = 1.84g (60%). ^1^HNMR (CDCl_3_): δ 11.05 (br s, 1H), 8.36 (s, 1H), 7.15 (d, 1H, *J* = 4.0Hz), 6.63 (br s, 1H), 5.10 (dd, 1H, *J* = 51.2, 13.6Hz), 4.91 (s, 0.5H), 4.71 (s, 0.5H), 4.61 (m, 1H), 3.98 (dd, 1H, *J* = 30.2, 12.4Hz), 3.67 (s, 3H), 3.20-3.51 (m, 4H); ^13^C NMR (CDCl_3_): {rotamer signals} {171.31, 170.94}, 157.30, {155.67, 155.17}, 152.21, 150.73, 121.37, 103.50, 101.35, 81.12, {55.92, 54.45}, 52.58, {46.97, 45.73}, {45.55, 44.92}, {41.81, 40.36}, 28.55; MS (ES+, M+1): 362.2.

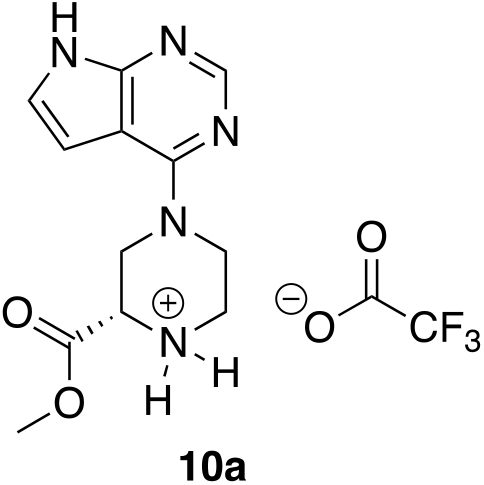

#### Methyl (*S*)-4-(*7H*-pyrrolo[2,3-*d*]pyrimidin-4-yl)piperazine-2-carboxylate •TFA (10a)

1-(*tert*-butyl) 2-methyl (*S*)-4-(*7H*-pyrrolo[2,3-*d*]pyrimidin-4-yl)piperazine-1, 2-dicarboxylate **9a** (1.642 g, 4.54 mmols), DCM (6 mL) and TFA (2.5 mL) were combined into a 100 mL round bottom flask and left stirring with no heat for 24hrs. TLC in a 100% EtOAc solvent system indicated that the reaction was complete. The DCM was removed by rotary evaporation. 5-10 mL of DCM and 10 mL of toluene were subsequently added to and evaporated via rotary evaporation from the round bottom containing the reaction mixture 5 times to extract the excess TFA. Dry yield (TFA salt) = 1.7g (quant). ^1^HNMR (DMSO-*d_6_*): δ 12.44 (br s, 1H), 8.42 (br s, 1H), 8.38 (s, 1H), 7.39 (dd, 1H, *J* = 4.0, 2.0Hz), 6.75 (d, 1H, *J* = 4.0Hz), 4.83 (dd, 1H, *J* = 15.2, 2.8Hz), 4.49 (m, 2H), 3.78 (s, 3H), 3.73 (m, 2H), 3.45 (m, 1H), 3.23 (m, 1H); ^13^C NMR (DMSO-*d_6_*): (TFA salt) 168.17, 167.14 (TFA), 158.27 (TFA, q, *J_C-F_* = 34.3Hz), 155.58, 151.15, 149.66, 122.88, 102.67, 100.52, 54.31, 54.08, 53.21, 44.30, 42.28; MS (ES+, M+1): 262.2.

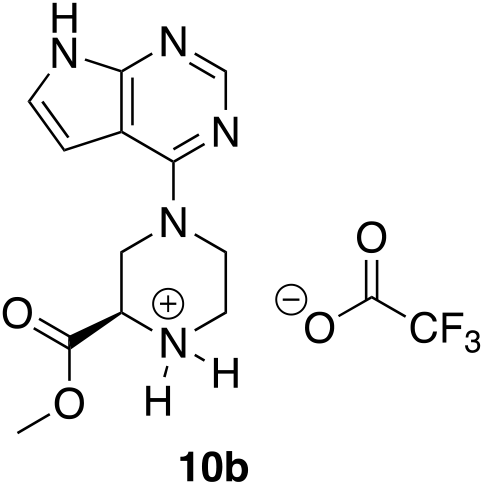

#### Methyl (*R*)-4-(*7H*-pyrrolo[2.3-*d*]pyrimidin-4-yl)piperazine-2-carboxylate •TFA (10b)

This compound was made in an identical manner to **10a** using **9b** as the starting material. Dry yield = 0.48g (quant). ^1^HNMR (DMSO-*d_6_*): δ 12.5 (br s, 1H), 8.42 (br s, 1H), 8.38 (s, 1H), 7.39 (dd, 1H, *J* = 4.0, 2.0Hz), 6.75 (d, 1H, *J* = 4.0Hz), 4.83 (dd, 1H, *J* = 15.2, 2.8Hz), 4.49 (m, 2H), 3.78 (s, 3H), 3.73 (m, 2H), 3.45 (m, 1H), 3.23 (m, 1H); ^13^C NMR (DMSO-*d_6_*): (TFA salt) 168.17, 167.14 (TFA), 158.27 (TFA, q, *J_C-F_* = 34.3Hz), 155.58, 151.15, 149.66, 122.88, 102.67, 100.52, 54.31, 54.08, 53.21, 44.30, 42.28; MS (ES+, M+1): 262.2.

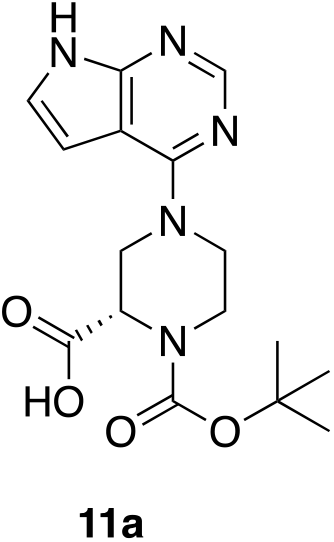

#### (*S*)-1-(*tert*-butoxycarbonyl)-4-(*7H*-pyrrolo[2,3-*d*]pyrimidin-4-yl)piperazine-2-carboxylic acid (11a)

Compound **9a** (0.5g, 1.38mmol), 1.25M aq. NaOH (4 mL, 5 mmol) and MeOH (4 mL) were combined into a 50 mL round bottom flask and left stirring below the boiling point overnight. The MeOH was removed via rotary evaporation. The remaining reaction mixture was transferred to a 125 mL separatory funnel and suspended in 5-10 mL of EtOAc. The layers were separated and the basic layer was acidified drop-wise with 3M HCl to 3-4 pH. The acidic layer was extracted twice with 10 mL of EtOAc. The combined organic layers were dried with solid MgSO_4_, vacuumed filtered and concentrated via rotary evaporation. Dry yield = 95mg (19%). ^1^HNMR (CDCl_3_): δ 11.91 (bs, 1H), 8.24 (s, 1H), 7.15 (s, 1H), 6.62 (s, 1H), 4.83 (s, 1H), 4.09 (m, 2H), 3.59 (m, 4H), 1.27 (s, 9H). ^13^C NMR (DMSO-*d_6_*): {172.00, 171.74}, {156.51, 156.43} {154.77, 154.51}, 151.87, 150.38, 121.74, 102.49, {100.58, 100.52}, {79.58, 79.40}, 55.20, 53.83, {45.55, 45.02}, {44.64, 44.34} 41.20, {27.99, 27.85}; MS (ES+, M+1): 348.2.

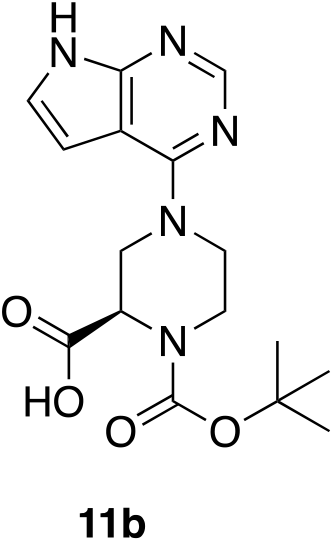

#### (*R*)-1-(tert-butoxycarbonyl)-4-(*7H*-pyrrolo[2,3-*d*]pyrimidin-4-yl)piperazine-2-carboxylic acid (11b)

This compound was made in an identical manner to compound **11a** using **10b** as the starting material. Dry yield = 212mg, 44.1%. ^1^HNMR (CDCl_3_): δ 11.91 (bs, 1H), 8.24 (s, 1H), 7.15 (s, 1H), 6.62 (s, 1H), 4.83 (s, 1H), 4.09 (m, 2H), 3.59 (m, 4H), 1.27 (s, 9H); ^13^C NMR (DMSO-*d_6_*): {172.00, 171.74}, {156.51, 156.43} {154.77, 154.51}, 151.87, 150.38, 121.74, 102.49, {100.58, 100.52}, {79.58, 79.40}, 55.20, 53.83, {45.55, 45.02}, {44.64, 44.34} 41.20, {27.99, 27.85}; MS (ES+, M+1): 348.2.

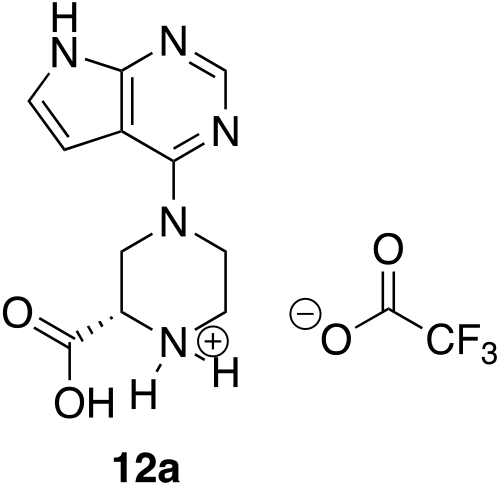

#### (*S*)-4-(*7H*-pyrrolo[2,3-*d*]pyrimidin-4-yl)piperazine-2-carboxylic acid (12a)

(*S*)-1-(*tert*-butoxycarbonyl)-4-(*7H*-pyrrolo[2,3-*d*]pyrimidin-4-yl)piperazine-2-carboxylic acid (28mg, 0.08mmol), DCM (1 mL) and TFA (1 mL) were combined into a 25 mL round bottom flask and left stirring with no heat for 1h. The DCM was removed via rotary evaporation. 5-10 mL of DCM were subsequently added to and evaporated via rotary evaporation from the round bottom containing the reaction mixture 7 times to extract the excess TFA. The product was characterized as the TFA salt. Dry yield = 33mg (quant). ^1^HNMR (DMSO-*d_6_*): δ 11.96 (bs, 1M), 9.60 (br s, 2H), 8.26 (s, 1H), 7.32 (dd, 1H, *J* = 4.0, 2.8Hz), 6.64 (dd,1H, *J* = 4.0, 2.8Hz), 4.80 (dd, 1H, *J* = 14.0, 2.8Hz), 4.43 (d, 1H, *J* = 14.0Hz), 4.31 (dd, 1H, *J* = 10.0, 3.6Hz), 3.64-3.77 (m, 2H), 3.40 (d, 1H, *J* = 12.8Hz), 3.16 (t, 1H, 10.0Hz); ^13^C NMR (DMSO-*d_6_*): 168.21, 158.31 (TFA, q, *J_C-F_* = 33.4Hz), 155.61, 151.19, 149.57, 122.90, 102.68, 100.54, 54.32, 44.31, 42.21, 42.01; MS (ES+, M+1): 248.1.

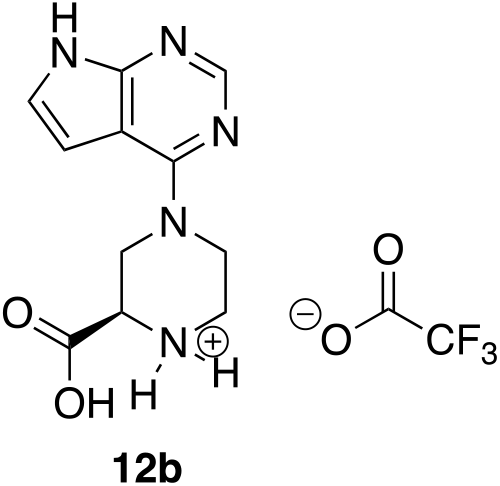

#### (*R*)-4-(*7H*-pyrrolo[2,3-*d*]pyrimidin-4-yl)piperazine-2-carboxylic acid (12b)

This compound was made in an identical manner to **12a** using **11b** as the starting material. Dry yield = 245mg (quant). 12.16 (bs, 1M), 9.60 (br s, 2H), 8.33 (s, 1H), 7.36 (d, 1H, *J* = 4.0Hz), 6.70 (d,1H, *J* = 4.0Hz), 4.82 (dd, 1H, *J* = 14.0, 2.8Hz), 4.45 (d, 1H, *J* = 14.0Hz), 4.33 (dd, 1H, *J* = 10.0, 3.6Hz) 3.64-3.77 (m, 2H), 3.42 (d, 1H, *J* = 12.8Hz), 3.19 (t, 1H, 10.0Hz); ^13^C NMR (DMSO-*d_6_*): 168.14, 158.23 (q, *J_C-F_* = 34.2Hz), 155.52, 150.31, 148.92, 123.13, 102.68, 100.87, 54.35, 44.44, 42.34, 42.00; MS (ES+, M+1): 248.1.

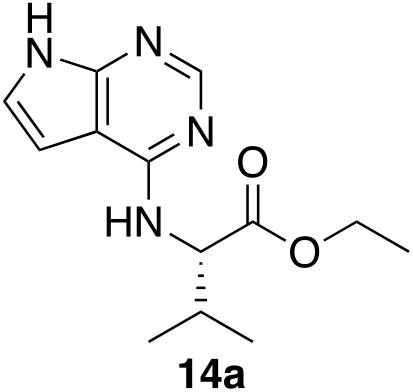

#### ethyl (*7H*-pyrrolo[2,3-*d*]pyrimidin-4-yl)-L-valinate (14a)

4-Chloro-*7H*-pyrrolo[2,3-*d*]-pyrimidine (0.500g, 3.26 mmol), L-Valine ethyl ester hydrochloride (0.712g, 3.92mmol), isopropanol (4 mL), and DIEA (1.71mL, 9.81mmol) were combined in a 25mL round bottom and left to stir under heat and water cooled condenser for 7 days. TLC showed that the reaction was complete, therefore the isopropanol was removed via rotary evaporator. EtOAc (10mL) was added to the reaction mixture, and poured into a separatory funnel. It was washed twice with 10% aqueous sodium bicarbonate and the organic layer was dried with MgSO_4_. The product was purified through a column chromatography on silica gel and was characterized as the desired compound. Dry yield: 0.170 g (20%). ^1^HNMR (CDCl_3_): δ 10.6 (s, 1H), 8.35 (s, 1H), 7.05 (d, 1H, *J* = 4.0Hz), 6.42 (d, 1H, *J* = 4.0Hz), 5.91 (d, 1H, *J* = 4.0Hz), 4.67 (q, 2H, *J* = 8.0Hz), 2.20 (m, 1H), 0.82-1.27 (m, 9H); ^13^C NMR (DMSO-*d_6_*): 172.61, 155.74, 150.93, 150.34, 120.98, 102.61, 99.10, 60.04, 58.85, 30.07, 19.20, 14.15; MS (ES+, M+1): 263.2.

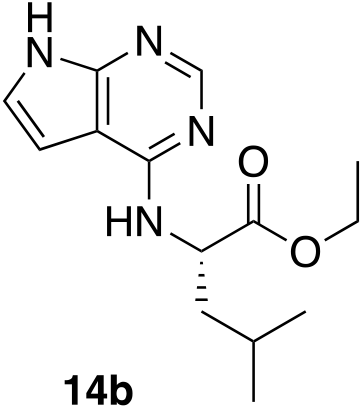

#### Ethyl (*7H*-pyrrolo[2,3-*d*]pyrimidine-4-yl)-*L*-leucinate (14b)

4-Chloro-7H-pyrrolo[2,3-*d*]-pyrimidine **3** (0.5g, 3.27mmol), L-Leucine-Ethyl-Ester-Hydrochloride (0.767g, 3.92mmol), DIEA (1.71ml, 9.81mmol), and isopropanol (4mL) were combined into a 25 mL round bottom flask and left to stir under heat, for approximately 48h. TLC indicated that the reaction was complete. The isopropanol was removed with the rotary evaporator and 10 mL of Ethyl Acetate was added to the reaction mixture. It was poured into a separatory funnel and washed with 5 mL of 10% NaHCO_3_. The organic layer was dried with MgSO_4_ and filtered into a tared round bottom flask. The product was purified via column chromatography (EtOAc). Dry yield = 0.31g (34%). ^1^HNMR (CDCl_3_): δ 8.27 (s, 1H), 7.02 (d, 1H,), 6.38 (s, 1H), 5.62 (s, 1H), 5.00 (s, 1H), 4.22 (m, 2H), 1.6-1.85 (m, 3H), 1.26 (t, 3H), 0.94 (d, 3H); ^13^C NMR (DMSO-*d_6_*): 173.64, 155.57, 151.01, 150.23, 121.02, 102.56, 98.79, 60.14, 51.35, 24.38, 22.81, 21.26, 14.06; MS (ES+, M+1): 277.1.

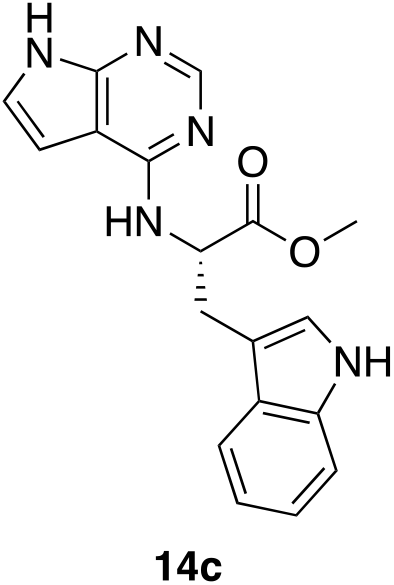

#### Methyl (*7H*-pyrrolo[2,3-*d*]pyrimidin-4-yl)-L-tryptophanate (14c)

methyl 4-Chloro-*7H*-pyrrolo[2,3-*d*]-pyrimidine **3** (0.500g, 3.27 mmol), L-Tryptophan Methyl Ester Hydrochloride (0.998g, 3.92mmol), isopropanol (4mL) and DIEA (1.71mL, 9.81mmol) were combined in a 25mL round bottom and left to stir under reflux for 5 days. TLC indicated that the reaction was complete, therefore the isopropanol was removed via rotary evaporator. 10mL of EtOAc was added to the reaction mixture, and poured into a separatory funnel. It was washed twice with 10% sodium bicarbonate (3 mL), and the organic layer dried with MgSO_4_. The product was purified through a column chromatography on silica gel. Dry yield=0.220g (20.0%). ^1^HNMR (DMSO-*d_6_*): δ 9.41 (s, 1H), 8.38 (s, 1H), 8.10 (s, 1H), 7.58 (d, 1H), 7.38 (d, 1H), 7.20 (m, 1H), 7.10 (t, 1H), 7.02 (m, 1H), 6.28 (s, 1H), 5.60 (s, 1H), 5.39 (m, 1H), 5.0 (m, 1H), 4.2 (m, 2H), 3.57 (s, 3H), 3.50 (m, 2H); ^13^C NMR (DMSO-*d_6_*): 173.58, 155.37, 151.01, 150.23, 136.06, 127.04, 123.69, 121.15, 120.93, 118.40, 118.00, 111.43, 110.06, 102.58, 98.69, 54.02, 51.72, 27.27; MS (ES+, M+1): 336.1.

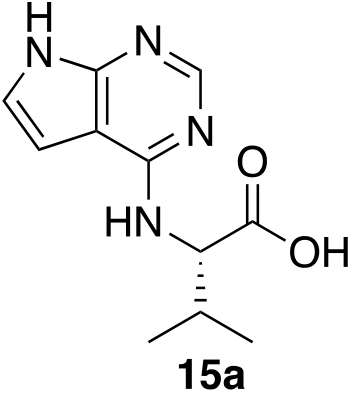

#### (7*H*-pyrrolo[2,3-*d*]pyrimidin-4-yl)-*L*-valine (15a)

Compound **14a** (0.150g, 0.50mmol), 1.25M NaOH (1.61mL, 2.02mmol) and 4mL of MeOH were combined in a 25mL round bottom flask and heated to reflux for 1h. TLC indicated that the reaction was complete. The MeOH was removed using the rotary evaporator. The product was acidified with 3M HCl and the solid was filtered using vacuum filtration. The dry yield was 0.113g (96%). ^1^HNMR (DMSO-*d_6_*): 11.49 (s, 1H), 8.05 (s, 1H), 7.29 (d, 1H, *J* = 4.0Hz), 7.07 (s, 1H), 6.77 (d, 1H, *J* = 4.0Hz), 4.61 (m, 1H), 2.19 (m, 1H), 1.00 (d, 3H, *J* = 6.4Hz), 0.97 (d, 3H, *J* = 6.4Hz); ^13^C NMR (DMSO-*d_6_*): 173.95, 155.90, 150.94, 150.28, 120.87, 102.61, 99.10, 58.56, 29.88, 19.29, 18.98; MS (ES+, M+1): 263.2.

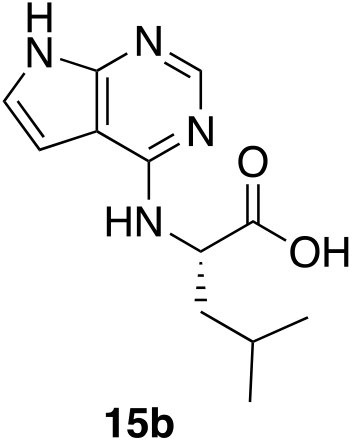

#### (*7H*-pyrrolo[2,3-*d*]pyrimidine-4-yl)-*L*-leucinate (15b)

The ethyl ester **14b** (0.270g, 0.77mmol), 1.25M NaOH (1.25ml, 3.08mmol) and 4mL of MeOH were combined in a 25mL round bottom flask and heat to boiling for 1h, using a water cooled condenser. TLC indicated that the reaction was complete. The MeOH was removed using the rotary evaporator. The product was acidified with 3M HCl and the solid was filtered using vacuum filtration. The dry yield was 0.082g (43%). ^1^HNMR (DMSO-*d_6_*): 11.5 (s, 1H), 8.05 (s, 1H), 7.47 (d, 1H), 7.06 (m, 1H), 6.64 (d, 1H), 4.85 (m, 1H), 1.76 (m, 2H), 1.62 (m, 1H), 0.92 (m, 6H); ^13^C NMR (DMSO-*d_6_*): 175.14, 155.78, 151.04, 150.17, 120.91, 102.60, 98.86, 51.22, 40.05, 24.46, 22.95, 21.25; MS (ES+, M+1): 249.1.

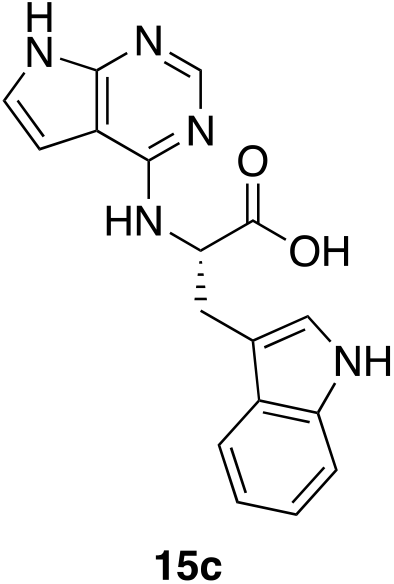

#### (*7H*-pyrrolo[2,3-*d*]pyrimidin-4-yl)-*L*-tryptophan (15c)

Compound **14c** (0.214g, 0.64mmol), 1.25M NaOH (2mL, 2.56mmol) and 4mL of MeOH were combined in a 25mL round bottom flask and heated to boiling for 1h. TLC indicated that the reaction was complete. The MeOH was removed using the rotary evaporator. The product was acidified with 3M HCl and the solid was filtered using vacuum filtration. The dry yield was 0.113g (55%). ^1^HNMR (DMSO-*d_6_*): 11.46 (s, 1H), 10.75 (s, 1H), 8.01 (s, 1H). 7.57 (d, 1H, *J* = 8.0Hz), 7.50 (s, 1H), 7.28 (d, 1H, *J* = 8.0Hz), 7.16 (s, 1H), 7.01 (m, 3H), 6.58 (s, 1H), 4.82 (m, 1H), 3.22 (m, 2H); ^13^CNMR (DMSO-*d_6_*): 174.52, 155.62, 151.06, 150.17, 136.05, 127.19, 123.60, 120.96, 120.83, 118.31, 118.19, 111.34, 110.60, 102.59, 98.70, 54.19, 27.23; MS (ES+, M+1): 322.1.

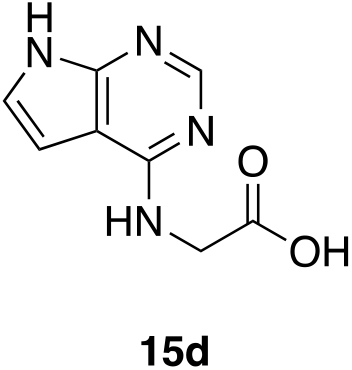

#### (*7H*-pyrrolo[2,3-*d*]pyrimidin-4-yl)glycine (15d)

4-Chloro-*7H*-pyrrolo[2,3-*d*]-pyrimidine (0.500g, 3.26 mmol, Glycine (0.294g, 3.92mmol), isopropanol (4 mL), and DIEA (1.71mL, 9.81mmol) were combined in a 25mL round bottom and left to stir under heat for 9 days. TLC showed that the reaction was complete, therefore the isopropanol was removed via rotary evaporator. EtOAc (10mL) was added to the reaction mixture, and poured into a separatory funnel. It was washed twice with 10% sodium bicarbonate (2 mL) and the aqueous layer was allowed to evaporate/concentrate overnight. The product eventually crashed out of the aqueous layer and was filtered off and characterized as the desired product. Dry yield = 0.263g (42%). ^1^HNMR (DMSO-*d_6_*). 11.5 (s, 1H), 8.05 (s, 1H), 7.58 (s, 1H), 7.07 (d, 1H), 6.53 (d, 1H), 4.05 (s, 2H); ^13^C NMR (DMSO-*d_6_*): 172.18, 155.82, 151.18, 150.16, 120.98, 102.62, 98.48, 42.44; MS (ES+, M+1): 193.1.

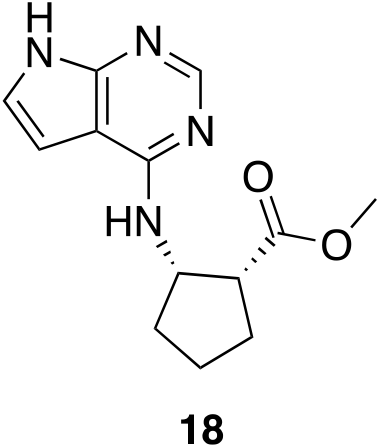

#### Methyl (1*R*, 2*S*)-2-((*7H*-pyrrolo[2,3-*d*]pyrimidin-4-yl)amino)cyclopentane-1-carboxylate (18)

A 25.0 mL round bottom was used to combine chloride **3** (0.401 g, 2.62 mmol), methyl (1*R*, 2*S*)-2-aminocyclohexane-1-carboxylate **17** (0.451 g, 3.15 mmol), DIEA (1.37 mL, or 7.86 mmol), and 3.0 mL of isopropanol. The reaction was heated below the bp for 1.5 weeks, TLC indicated that the reaction was still incomplete (EtOAc). The reaction was concentrated on the rotary evaporator to remove the isopropyl alcohol. 10 mL of EtOAc was added to the round bottom. The organic layer was washed with 10% NaHCO_3_ and the organic layer was dried with MgSO_4_, filtered off using vacuum filtration and was allowed to air dry. Column chromatography was conducted (80/20% EtOAc/Hexanes → 100% EtOAc→95/5 EtOAc/MeOH). Dry yield = 0.051g, 12.7% yield. ^1^HNMR (CDCl_3_): δ 11.78 (s, 1H), 8.32 (s, 1H), 7.07 (d, 1H, *J* = 4.0Hz), 6.39 (d, 1H, *J* = 4.0Hz), 5.58 (s, 1H), 4.82 (m, 1H), 3.63 (s, 3H), 2.80 (m, 1H), 2.32 (m, 1H), 1.60-2.05 (m, 5H); ^13^C NMR (DMSO-*d_6_*): 175.26, 155.59, 151.23, 150.13, 120.75, 102.43, 98.54, 55.00, 51.36, 49.61, 32.62, 28.77, 23.13; MS (ES+, M+1): 261.1.

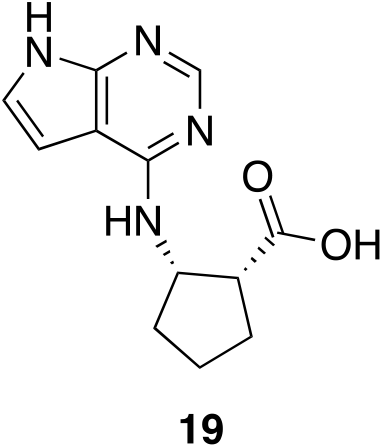

#### (1*R*, 2*S*)-2-((*7H*-pyrrolo[2,3-*d*]pyrimidin-4-yl)amino)cyclopentane-1-carboxylic acid (19)

A 25.0 mL round bottom was used to combine the ester **18** (0.033 g, 0.127 mmol) and 10 mL of 3M HCl. The reaction was heated to reflux for 3 hours. Rotary evaporation was then used to remove the aqueous liquid. The solid was triturated with cold water and filtered to afford the product. Dry yield = 0.021g, 63.6%. (DMSO-*d_6_*): δ 12.79 (s, 1H), 9.98 (s, 1H), 8.30 (s, 1H), 7.37 (s, 1H), 7.15 (s, 1H), 4.65 (m, 1H), 3.19 (m, 1H), 2.06-1.64 (m, 6H); ^13^C NMR (DMSO-*d_6_*): 175.00, 150.95, 150.01, 142.31, 124.19, 102.35, 101.67, 55.77, 48.53, 32.20, 28.04, 22.76; MS (ES+, M+1): 247.1.

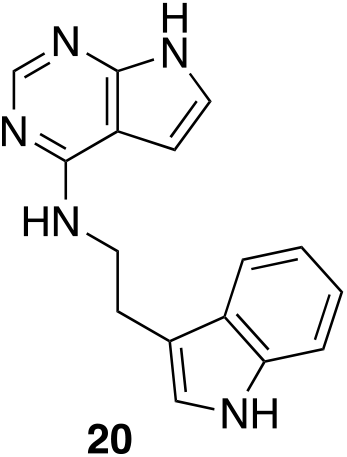

#### *N*-(2-(1*H*-indol-3-yl)ethyl)-*7H*-pyrrolo[2,3-*d*]pyrimidin-4-amine (20)

4-chloro-*7H*-pyrrolo[2,3-*d*]pyrimidine **3** (201 mg, 1.31 mmol), tryptamine (253 mg, 1.58 mmol), isopropanol (3.5 mL), and DIEA (0.35 mL, 2.01 mmol) and a stir bar were placed into a 25 mL round bottom flask and allowed to stir at 70 °C overnight. TLC provided evidence of a complete reaction (1:7 MeOH-EtOAc). The isopropanol was removed via rotary evaporator, and to the remaining brown oil was added 2mL of EtOAc and 2 mL of dH_2_O. Approximately 1 hour later, a tan solid began forming between the EtOAc-diH_2_O interface. With a small application of heat more solid began forming. Once no more solid formed the mixture was vacuum filtered to collect the solid. The solid was then placed in a vacuum oven to dry. Dry yield 126 mg (34.7% yield). ^1^HNMR (DMSO-*d_6_*): δ 11.45 (s, 1H), 10.80 (s, 1H), 8.12 (s, 1H), 7.59 (d, 1H), 7.51 (m, 1H), 7.32 (d, 1H), 7.17 (d, 1H), 7.01 (m, 3H), 6.51 (dd, 1H), 3.73 (m, 2H), 2.99 (m, 2H); ^13^C NMR (DMSO-*d_6_*): 156.05, 151.51, 150.06, 136.23, 127.32, 122.57, 120.88, 120.62, 118.36, 118.20, 112.10, 111.33, 102.52, 98.54, 40.86, 25.35; MS (ES+, M+1): 278.1.

